# Pre-clinical validation of a novel AAV-mediated gene therapy for *KCNV2* retinopathy improves visual function in a mouse model and expression in patient organoids

**DOI:** 10.64898/2025.12.09.692426

**Authors:** Rabab Rashwan, Paula I Fuller-Carter, Xin Ru Lim, Alicia A Brunet, Annie L Miller, Yashvi Bhatt, Rebekah James, Denise Anderson, Valentina Voigt, Joao A Paulo, Mehdi Mirzaei, Melissa M Mangala, Emilie O Wong, Robyn V Jamieson, Anai Gonzalez-Cordero, David M Hunt, Livia S Carvalho

## Abstract

Voltage-gated (Kv) potassium channels are critical for neuronal physiology, and their dysfunction can lead to serious consequences. For example, mutations in the silent modulatory Kv8.2 subunit are known to cause irreversible inherited blindness (*KCNV2* retinopathy). This is a currently incurable condition that causes lifelong visual loss, reduced visual acuity, photoaversion, night blindness and abnormal colour vision, alongside a distinctive supernormal electrophysiological (ERG) retinal response to light. In this study, we demonstrate that AAV-mediated gene replacement therapy delivering a codon-optimised human *KCNV2* gene subretinally into Kv8.2 knock-out mice significantly restores retinal function. Treated mice exhibited improved ERG responses and correct expression of KCNV2 and its encoded Kv8.2 protein in photoreceptors. Recovery of visually guided scotopic and photopic optomotor responses to wildtype levels was achieved at lower vector doses, highlighting dose-dependent efficacy. Furthermore, treatment of human retinal organoids derived from a KCNV2 patient iPSC line resulted in substantial Kv8.2 protein rescue. This work provides the first preclinical proof-of-concept for the safety and therapeutic potential of gene therapy for KCNV2 retinopathy, laying a strong foundation for future clinical trials.

## Introduction

*KCNV2* retinopathy or Cone dystrophy with supernormal rod response [CDSRR] (Retinal cone dystrophy 3B, OMIM 610356), is a rare, recessive and inherited retinopathy with an estimated prevalence of around 1/800,000 (1, 2). It is characterised by poor visual acuity (due to central scotoma), photophobia, severe colour vision deficits, with nystagmus and strabismus occasionally present (3–6). Visual difficulties begin in early childhood with visual acuity of 20/100 or worse by the second decade of life. Patients will later develop night blindness, and most will also develop significant myopia (7–9). The fundus appears normal in some patients but foveal or parafoveal atrophy, a macular bull’s eye, hyperfluorescence anomalies, and a generalised fine pigmentary retinopathy have been reported, and there may be some temporal pallor in the optic nerve (9).

Clinical symptoms are restricted to visual loss with no other tissues or organs affected. Critically, no specific treatment is available to either reduce or prevent the progression of visual loss, leaving patients with a poor prognosis and declining quality of life as they age (10, 11). KCNV2 retinopathy is characterised by an altered electroretinography (ERG) response; there are reduced and delayed dark adapted (rod photoreceptor specific, or scotopic) a- and b-waves, with the latter uniquely switching to an enhanced response with supernormal amplitude at higher light intensities. This unusual ERG is diagnostic for the disorder and can be used to monitor its progression. In contrast, cone photoreceptor specific (photopic) ERGs are markedly reduced and delayed over the entire range of light intensities (12–16).

Genetic lesions in the *KCNV2* gene that encodes the Kv8.2 voltage-gated potassium (K^+^) channel subunit are the cause behind this condition (5, 17), with more than 263 pathogenic and likely pathogenic variants(18) now identified, including missense and nonsense mutations, intragenic deletions, and out-of-frame insertions (3–5, 10, 12–16, 19–23). Although the different variants have been shown to affect Kv8.2 differently, with some generating non-conducting channels, whereas others prevent channel formation altogether (24), it would appear that all variants result in the absence of functional channels containing the Kv8.2 subunit.

Kv8.2 is a member of a group of “modifier/silent” channel proteins that do not form channels by themselves but require a cognate partner; for Kv8.2, this is Kv2.1 (encoded by *KCNB1*) (24, 25), a member of the Shab family of subunits that generate delayed rectifier currents important in the regulation of the rate of repolarisation of action potentials. In the eye, both the Kv8.2 and Kv2.1 subunits are located exclusively on the cytoplasmic membrane of the inner segments of cone and rod photoreceptor cells, located in the retinal tissue in the eye and responsible for initiating the light transduction cascade of the visual response (26).

Two recent studies carried out in our laboratory characterised a *Kcnv2* knock-out (KO) mouse model of the human disorder, the C57BL/6N*_Kcnv2^tm1^* mouse (referred to as Kv8.2 KO hereafter)(27, 28). Detailed analysis of the Kv8.2 KO mice(27) shows that the ERG is severely depressed at low light flash intensities but then switches to an enhanced response at high light flash intensities, in parallel to the response in human patients. The overall thickness of the photoreceptor layer in Kv8.2 KO mice showed significant thinning of about 40% compared to wildtype, although the photoreceptor outer segments appear to be intact. Cone photoreceptors showed only a 20% loss by 6 months of age, indicating that the thinning is largely the result of rod photoreceptor loss. There is a significant increase in retinal cell death at 1 month of age compared to WT, and this difference remains significant at 3 and 6 months, although the overall frequency is lower. The damage to the retina from the absence of Kv8.2 subunits is therefore relatively mild, with the majority of photoreceptors (both rods and cones) retained in the retina. This is paralleled by a recent pupillometry study of inner retinal function in CDSRR patients which indicated that this function may be largely preserved. These findings, together with the recessive mode of inheritance and slow progression of the disorder (1, 2, 7, 29), make it a good candidate for a one-off viral-based gene therapy (gene supplementation) treatment (9, 30). In addition, the unique ERG presentation allows for an accurate and early diagnosis of the disorder besides also providing a reliable visual function outcome measure of treatment efficacy.

Several animal models of inherited photoreceptor degeneration have undergone successful treatment by gene supplementation therapy, and to date visual rescue has been achieved on morphological, functional and behavioural levels (31). There are 65 ongoing or completed clinical trials using AAV delivery systems as a means of correcting genetic faults for different types of inherited retinal disorder (clinicaltrials.gov). In parallel, we explored the utility of human retinal models derived from patient induced pluripotent stem cells (iPSCs). Organoids can provide a relatively quick and cost-effective readout of gene augmentation efficacy showing that a viral vector construct can effectively work on human cells, in particularly, human photoreceptor cells (32).

In this study, we report the outcomes of a gene therapy treatment approach in the Kv8.2 KO mouse using AAV vectors to deliver a codon-optimised version of the human *KCNV2*, under the control of the human *Rhodopsin Kinase (RK)* promoter, to the photoreceptor cells in the retina. Treatment efficacy has been analysed by several ERG measures, gene and protein expression and vision dependent behavioural testing. We show for the first time, that the unusual ERG phenotype presented by the absence of Kv8.2, can be corrected with gene replacement therapy as early as 4 weeks post-treatment. We also established that despite the relatively slow progression of the disease, treatment efficacy is time-dependent, but recovery of some visual outcomes can still be achieved, even when the treatment is delivered to 6 months old mice. This successful rescue was corroborated by a significant overexpression of around 50% of the Kv8.2 protein in a patient iPSC-derived retinal organoid model. Overall, this study confirms the high suitability of our AAV-based *KCNV2* gene therapy as a strong candidate for future clinical trials.

## Results

### Low dose treatment significantly impacts the scotopic positive b-wave ERG

Previous studies by us and others (26, 28, 33–35) have confirmed that both cone and rod photoreceptors express heteromeric Kv2.1/Kv8.2 channels in their inner segments. This confirms the long-held belief that the *I_kx_* potassium current driving the dark resting potential of photoreceptors is in turn driven by voltage-gated potassium channels (36–38). Given the importance of Kv channels in regulating ionic flux and thereby the ability of photoreceptors to respond to light, it is not surprising that loss of functional Kv8.2 subunits generates an abnormal electrophysiological response in the retina to light. Furthermore, the expression of these channels needs to be tightly controlled and cell-specific, as overexpression could also generate dysfunction and under expression might not be sufficient to provide physiological recovery. To this end, it was essential that our vectors contained a promoter that drives transgene expression exclusively where *KCNV2* is expressed in the rod and cone photoreceptor cells. We selected the short *RK* promoter (39) which has been previously shown to restrict expression to rod and cone photoreceptors, and is currently in use in AAV vectors for clinical trials for the treatment of IRDs (40). The AAV serotypes used in this study, AAV8 and Anc80, were selected based on their affinity for photoreceptor tropism, a strong expression pattern and, in the case of AAV8, its utilisation in clinical trials for IRDs(40–46). Our first step was to evaluate the impact of subretinal injections on our main outcome measure, the electroretinogram (ERG). Neither sham (1µl PBS) or control AAV vector (Anc80.GFP and AAV8.GFP) injections had any effect on ERG amplitudes up to 12 weeks post-injection in wild type (WT) mice treated at P30 (Supplementary Fig. 1S-A-B).

Given the importance of keeping K^+^ flux well regulated, we initially evaluated the effect of a low dose of 1x10^9^ vector genomes (1e9 vg) for both our AAV8-KV and Anc80-KV vectors on the ERG response in Kv8.2 KO mice. We have previously demonstrated that the dark-adapted (scotopic) ERG response of Kv8.2 KO mice mimics the human phenotype with an abnormally increased scotopic b-wave response at high light intensities, while the a-wave is significantly reduced at all intensities(28, 33). At this low dose, the treatment with either vector was capable of significantly reducing the scotopic positive b-wave, but no improvement was observed in the scotopic a-wave (Supplementary Fig. 1S-C-D). On the photopic ERG response, neither a- nor b-waves were significantly improved after treatment (Supplementary Fig. 1S-E-F).

### Early and late treatment with AAV8 and Anc80 vectors show significant improvement in scotopic vision

Given that the vector dose of 1e9 vg did not have any negative impact on photoreceptor function in the Kv8.2 KO mice and was capable of driving Kv8.2 expression in the inner segments, the physiologically correct location for photoreceptor expression (data not shown), the vector dose was increased to 3e9 vg for Anc80-KV and to 3e9 vg (AAV8.KV) and 5e9 vg (AAV8.KV-HD) for AAV8 vectors. Kv8.2 KO mice treated at an early disease stage (P30) with either AAV8 or Anc80 (Fig. 1A) showed a significant improvement in a-wave amplitude compared to untreated Kv8.2 KO, at the increasing light intensities tested (*p* < 0.03; Fig. 1B). This improvement was already present at 4 weeks post-treatment for AAV8.KV-HD and remained present at the 8- and 12-week time points analysed (*p* < 0.0002; Fig. 1C). Interestingly, a significant improvement in a-wave amplitude was only observed in the Anc80 (3e9 vg) and AAV8 HD (5e9 vg) treated animals, but not in the 3e9 vg AAV8 treated group. All three vectors/doses showed a significant decrease in the positive b-wave amplitude from the abnormal amplitude present in untreated Kv8.2 KO mice at all time points (*p* < 0.0001; Fig. 1D-E). Furthermore, the positive b-wave amplitudes in all three treated groups returned to the WT level at low light intensities (*p* > 0.1; Fig. 1D).

**Figure 1.**
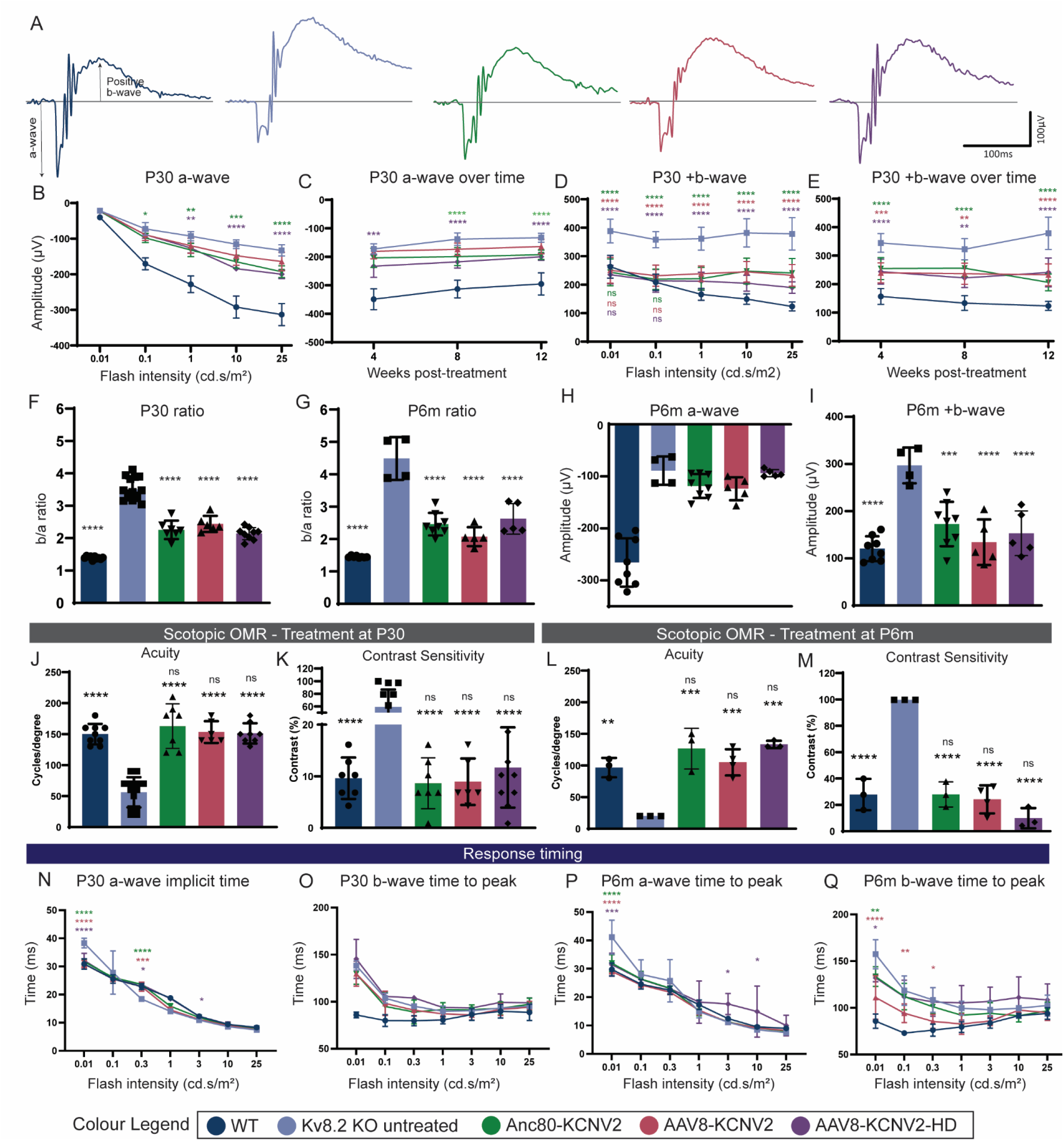
Improved dark-adapted ERG and optomotor responses after subretinal treatment with AAV-KCNV2 at 2 different ages. **(A)** Representative flash scotopic ERG traces at 25 cd.s/m^2^ recorded at 12 weeks post-treatment for age-matched wild type (WT) and Kv8.2 KO untreated mice alongside Kv8.2 KO treated mice at 30 days of age (P30) with AAV8.KCNV2 (1e9 vg), Anc80-KCNV2 (1e9 vg) and AAV8-KCNV2-HD (3e9 vg). **(B-E)** Quantification of scotopic ERG waves from P30 treated groups and age-matched controls at different intensities and overtime: (B) a-wave amplitudes at different flash intensities; (C) a-wave amplitudes over time at 4, 8 and 12 weeks; (D) positive b-wave amplitudes at different flash intensities; (E) positive b-wave amplitudes over time at 4, 8 and 12 weeks. **(F-G)** Quantification of b-wave to a-wave scotopic ratio at 25 cd.s/m2 for animals treated at P30 (F) and P6m (G). **(H-I)** Quantification of scotopic a-wave (H) and positive b-wave (I) recorded at 25 cd.s/m^2^ at 4 weeks post-treatment in Kv8.2 KO mice treated at 6 months of age (P6m) alongside age-matched WT and untreated Kv8.2 KO mice. **(J-M)** Dark-adapted optomotor reflex responses (OMR) for acuity and contrast sensitivity readouts for Kv8.2 KO mice treated at P30 and P6m alongside age-matched WT and Kv8.2 KO untreated controls. **(N-Q)** Implicit times to peak of a- and b-wave responses in animals treated at P30 (N-O) and P6m (P-Q). Numbers (n) are eyes from ≥3 biological replicates: (P30) WT n=8-10; Kv8.2 KO untreated n=8; Anc80-KCNV2 n=3-8; AAV8-KCNV2 n=4-5; AAV8-KCNV2 HD n=7-10; (P6m) WT n=3-8; Kv8.2 KO untreated n=3-4; Anc80-KCNV2 n=6-7; AAV8-KCNV2 n=5-6; AAV8-KCNV2 HD n=3-5. Bar graphs are averages ± SEM. Statistical analysis: two-way ANOVA followed by Tukey’s post hoc analysis. Significance is shown compared to Kv8.2 KO untreated: *p<0.03; **p<0.003; ***p<0.0002; ****p<0.0001. ns indicates non-significant compared to WT.

In the normal ERG response, since it is the electrical activity that is displayed as the a-wave that provides for activation of bipolar cells and the generation of the b-wave, there is generally a direct correlation between a-wave and b-wave amplitudes (47). In a normal mouse retina, the ratio between a- and b-wave amplitudes is around 1.5 (Fig 1F-G) whereas in the absence of Kv8.2 subunits in the KO mice, the functional link between the a- and b-wave amplitudes is lost, resulting in a significantly higher ratio (∼2.5 fold) (Fig. 1F-G). Our gene therapy treatment significantly improved this ratio at both doses and with both vectors (∼30-39% reduction, *p* < 0.0001; Fig. 1F), indicating that replacing the activity of the defective *KCNV2* gene with a normal copy can restore the relationship between activation of photoreceptors and downstream inner retinal responses from bipolar cells. Also, the ∼30% reduction in the b/a ratio correlates with the average area of the retina targeted with a subretinal injection (25-30% total retina area). This reduction in the b/a-wave ratio is even more pronounced in animals treated at later stages of disease (6 months of age), when the ratio difference between WT and untreated Kv8.2 KO mice is even larger due to the further degeneration of the a-wave response (∼3 fold; Fig. 1G). At 4 weeks post-treatment, the ratios for the P6m-treated groups have been reduced by 45%, 54% and 41.5% for Anc80.KV, AAV8.KV and AAV8.KV-HD, respectively (*p* < 0.0001; Fig. 1G). This reduction is substantial, considering that treatment at P6m was unable to significantly improve the a-wave, but was still capable of reducing the abnormal b-wave amplitude in animals treated with both vectors and at all three doses (*p* < 0.0002; Fig. 1H-I). In addition, treatment with either vector resulted in the full recovery of the visually guided optomotor reflex response (OMR), and this was independent of age at treatment and dose (Fig. 1J-M). Both scotopic contrast sensitivity and acuity responses were indistinguishable from WT (*p* > 0.09) and significantly improved from untreated (*p* < 0.002).

Time to peak (implicit time) of both a- and b-wave responses was also analysed and shown in Fig 1 panels N to Q. Treatment at both P30 or P6m, and with any of the three vectors, normalised the timing of the a-wave at low light intensities (Fig 1N,P), when the untreated retina is significantly delayed compared to wildtype (*p*<0.0001). The b-wave timing was however, not affected in the P30 treated retinas (Fig 1O), with treated responses remaining non-significant compared to untreated (*p*>0.2977). Interestingly, the b-wave implicit timing in the P6m treated retinas (Fig 1Q) was significantly reduced from untreated with all three vectors, probably due to the continued degeneration in the untreated retinas.

### Photopic response improvement is dependent on age, vector serotype and dose

*KCNV2* retinopathy is clinically classified as a cone-rod dystrophy, as its effect on the cone photoreceptors tends to be more severe and have an earlier onset than the impact on the rod system (3–5, 48, 49). This same pattern of degeneration is also seen in our Kv8.2 KO mouse model, where the loss of cone function (photopic) declines faster than rod function, despite a relatively modest loss of cone cells over time (28, 33). Figure 2 shows that our treatment results in an increase in the photopic ERG and light-adapted optomotor responses of Kv8.2 KO animals, indicating an improvement in the physiological response of cones (Fig. 2A,F - representative traces for the photopic flash and flicker responses of P30 treated mice), although a statistically significant recovery of cone function is less clear due to the naturally smaller amplitude of cone responses (Fig. 2B-E, G-H). What these results demonstrate, however, is that a higher dose is needed for a significantly improved a-wave response, and that a medium dose of the Anc80.KV vector can provide similar improvements in b-wave responses to the AAV8 vector at the higher dose (Fig. 2D-E). This was also confirmed by the cone flicker responses (Fig. 2G-H) where the best improvement was seen in the AAV8.KV-HD group. In contrast, however, to the scotopic ERG responses, no improvement was observed in the 6 months old-treated animals in their photopic flash or flicker responses, indicating that recovery of cone responses is more time-sensitive compared to rods.

**Figure 2.**
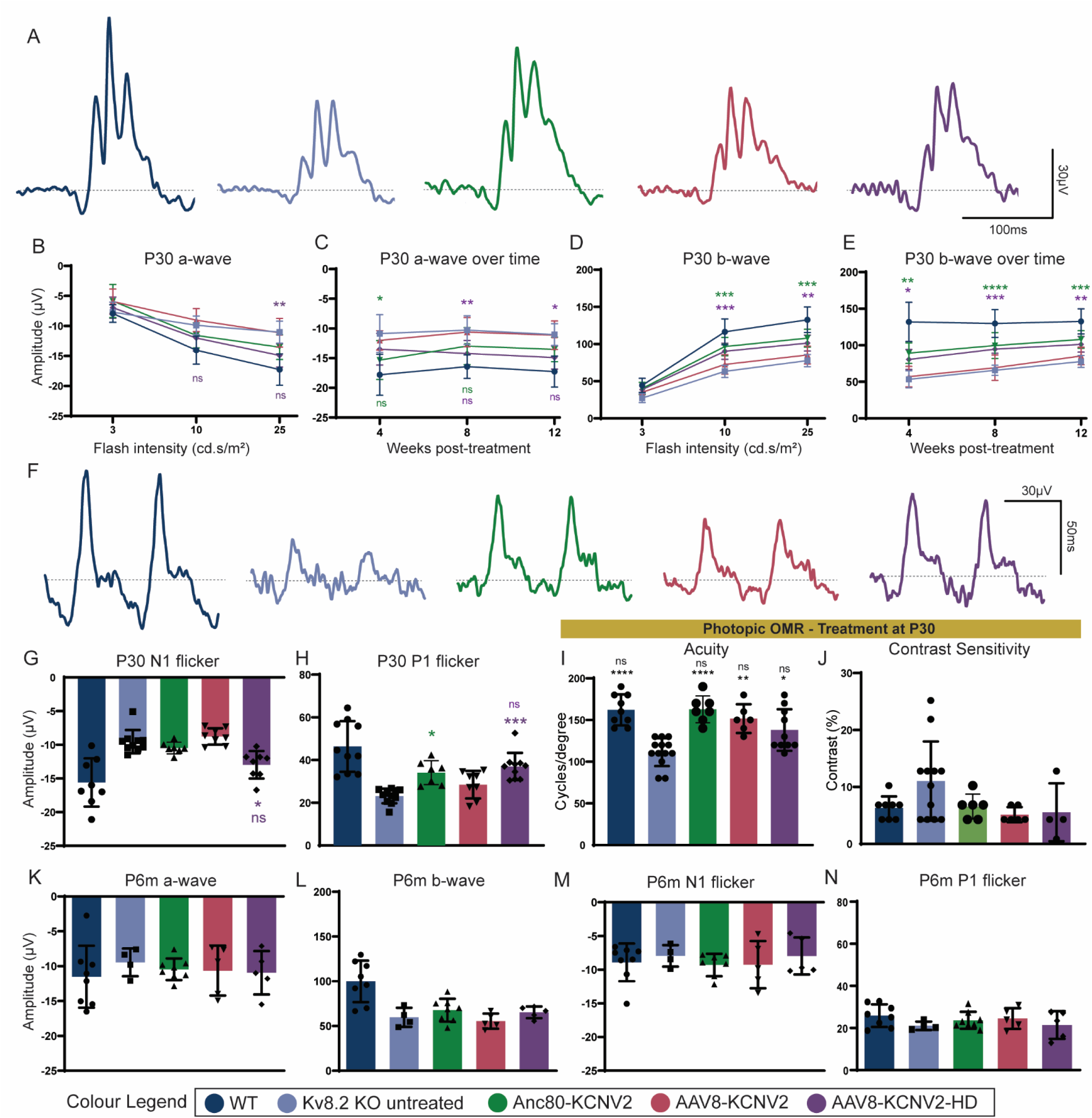
Light-adapted ERG and optomotor responses improve in mice treated at P30 but not P6m. **(A)** Representative flash photopic ERG traces at 25 cd.s/m^2^ recorded at 12 weeks post-treatment for age-matched wild type (WT) and Kv8.2 KO untreated mice alongside Kv8.2 KO treated mice at 30 days of age (P30) with AAV8.KCNV2 (1e9 vg), Anc80-KCNV2 (1e9 vg) and AAV8-KCNV2-HD (3e9 vg). **(B-E)** Quantification of photopic a- and b-waves from P30 treated groups and age-matched controls at different intensities and overtime: (B) a-wave amplitudes at different flash intensities; (C) a-wave amplitudes over time at 4, 8 and 12 weeks; (D) b-wave amplitudes at different flash intensities; (E) b-wave amplitudes over time at 4, 8 and 12 weeks. **(F)** Representative flicker ERG traces at 10 Hz recorded at 12 weeks post-treatment for age-matched wild type (WT) and Kv8.2 KO untreated mice alongside Kv8.2 KO treated mice at 30 days of age (P30) with AAV8.KCNV2 (1e9 vg), Anc80-KCNV2 (1e9 vg) and AAV8-KCNV2-HD (3e9 vg). **(G-H)** Quantification of P30 N1 and P1 flicker amplitudes at 12 weeks post-treatment. **(I-J)** Light-adapted optomotor responses for acuity and contrast sensitivity at 12 weeks post-treatment from mice treated at P30. **(K-N)** Quantification of photopic ERG responses at 4 weeks post-treatment from mice treated at P6m: (K) a-wave; (L) b-wave; (M) N1 flicker; (N) P1 flicker. Numbers (n) are eyes from ≥3 biological replicates: (P30) WT n=8-10; Kv8.2 KO untreated n=8-11; Anc80-KCNV2 n=6-8; AAV8-KCNV2 n=6-; AAV8-KCNV2 HD n=4-10; (P6m) WT n=8; Kv8.2 KO untreated n=4; Anc80-KCNV2 n=8; AAV8-KCNV2 n=4; AAV8-KCNV2 HD n=5. Bar graphs are averages ± SEM. Statistical analysis: two-way ANOVA followed by Tukey’s post hoc analysis. Significance is shown compared to Kv8.2 KO untreated mice: *p=0.02; **p<0.008; ***p<0.0007; ****p<0.0001. ns indicates non significance compared to WT.

### Electrophysiological treatment effects extend to other retinal cells beyond photoreceptors

The oscillatory potentials (OPs) are the 4-6 wavelets (OP1-6) found on the ascending limb of the ERG b-wave (50, 51). The current International Society for Clinical Electrophysiology of Vision (ISCEV) 2022 standards follow this definition of OPs, assigning their origins to inner retinal activity, specifically amacrine and retinal ganglion cells. Both rod and cone-mediated pathways are said to contribute to the generation of the OPs, with their origins being independent of the a- and b-waves, but dependent on the light stimuli and the state of adaptation of the retina (52–55). The absence of Kv8.2 subunits generates a very clear delay in the peak of OP1, which consequently shifts and delays both OP2 and OP3 (Fig. 2A). Our treatment was able to revert this timing delay of OP1-3 back to wild type, independent of the dose, vector or age at treatment (Fig. 3C-D), suggesting a wider range of beneficial effects beyond the directly affected and treated photoreceptor cells.

**Figure 3.**
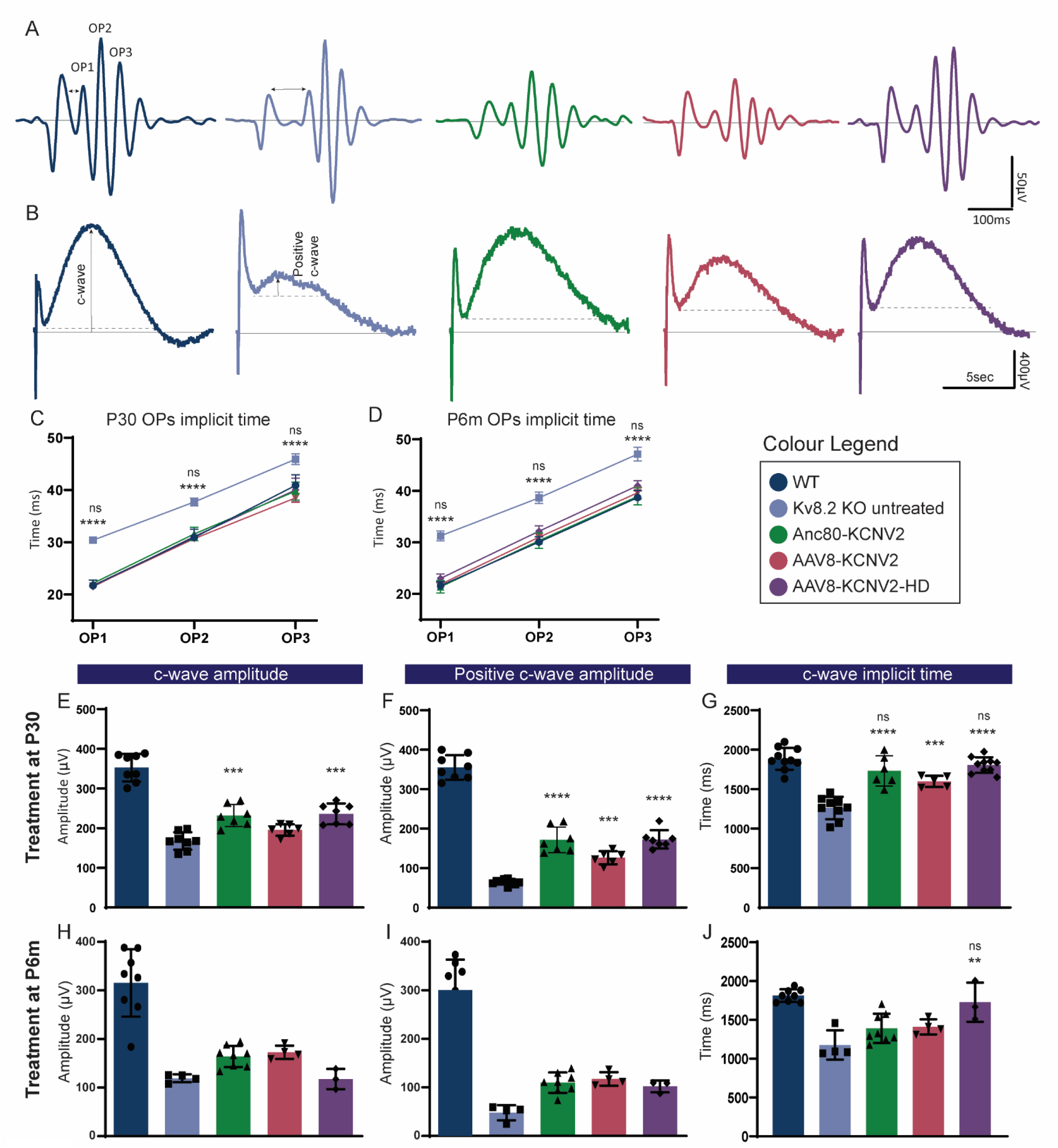
Treatment improves oscillatory potential timing and RPE-driven c-wave ERG response. **(A)** Oscillatory potentials (OPs) representative traces subtracted from the scotopic full flash response at 25 cd.s/m^2^ for WT, Kv8.2 KO untreated and Kv8.2 KO mice treated at P30 with Anc80-KCNV2, AAV8-KCNV2 and AAV8-KCNV2 HD. Arrows show the delay of time to peak of the Kv8.2 KO untreated OPs. **(B)** Representative c-wave traces for the same groups as in (A) and treated at P30. Dotted lines indicate the baseline used for the quantification of the positive c-wave amplitudes (arrows). **(C)** Quantification of OP1-3 amplitudes for WT, Kv8.2 KO untreated and Kv8.2 KO treated mice at P30 with Anc80-KCNV2, AAV8-KCNV2 and AAV8-KCNV2 HD. **(D)** OP1-3 implicit time quantification for all groups. WT n=10; Uninjected Kv8.2 KO n=12; Anc80-KCNV2 n=7; AAV8-KCNV2 n=6; AAV8-KCNV2 HD n=12. Line graphs are averages ± SEM. Statistical analysis: one-way ANOVA followed by Tukey’s post hoc analysis. ****p<0.0001. ns indicates non-significant compared to WT. **(E-G)** Quantification of ERG c-wave responses from all treated groups at P30 compared to wild type and untreated Kv8.2 KO mice showing amplitude measured from original baseline (E), amplitude measured from end of b-wave baseline (F), and implicit time to peak (G). Numbers (n) are eyes from ≥ 3 biological replicates: WT n=8-10; Kv8.2 KO untreated n=8-10; Anc80-KCNV2 n=6-7; AAV8-KCNV2 n=5-6; AAV8-KCNV2 HD n=7-10. Bar graphs are averages ± SEM. Statistical analysis: two-way ANOVA followed by Tukey ’ s post hoc analysis. Significance is shown compared to Kv8.2 KO untreated mice: **p=0.003; ***p=0.0006; ****p<0.0001. ns indicates non significance compared to WT.

Our previous study (35) showed that Kv8.2 KO mice display a reduction in the c-wave amplitude. The c-wave is believed to be generated by light-induced extracellular K^+^ uptake by the retinal pigment epithelium and Müller glia cells, which causes these cells to hyperpolarise (47). An altered K^+^ flux in photoreceptors due to the absence of Kv8.2 subunits will consequently impact the generation of the c-wave, which can also be used as an additional outcome measure of the efficacy of our therapeutic product. For animals treated at P30, improvements in c-wave amplitudes were more pronounced in the Anc80.KV 3e9 and AAV8.KV-HD 5e9 groups (Fig. 3E-F; *p*<0.0006), while an improvement in the c-wave time to peak (implicit time) was seen in all three groups (Fig. 3G; *p*<0.0006). When receiving treatment at 6 months of age, the only significant difference was observed in the implicit timing of the AAV8.KV-HD 5e9 group (*p*=0.003), and while a trend was observed in higher c-wave amplitudes, it was not significant (Fig. 3H-I). The improvement on c-wave amplitude and timing seen after treatment could indicate improved extracellular K^+^ flux in the retina due to the restoration of Kv channel function.

### Long-term restoration of visual response after treatment

As photoreceptor cells are post-mitotic and therefore do not divide, their loss in conditions such as inherited retinal diseases is irreversible. However, therein lies the advantage of an AAV-based gene replacement therapy as a one-off treatment. Figure 4 shows the still significant long-term effect of our AAV8.KV-HD gene therapy on scotopic visual function at 15 months post-treatment. Despite an amplitude decrease in treatment effect compared to the earlier timepoints, we still observed a significant improvement in the scotopic a- and b-wave response and b/a ratio (Fig 4A-C). Interestingly, the timing of the OPs was also significantly corrected after treatment back to wildtype levels (Fig 4D), as seen in the earlier timepoints evaluated. However, the significant improvement seen in earlier timepoints on the photopic a- and b-wave and c-wave was lost at 15 months (Fig 4E-H).

**Figure 4.**
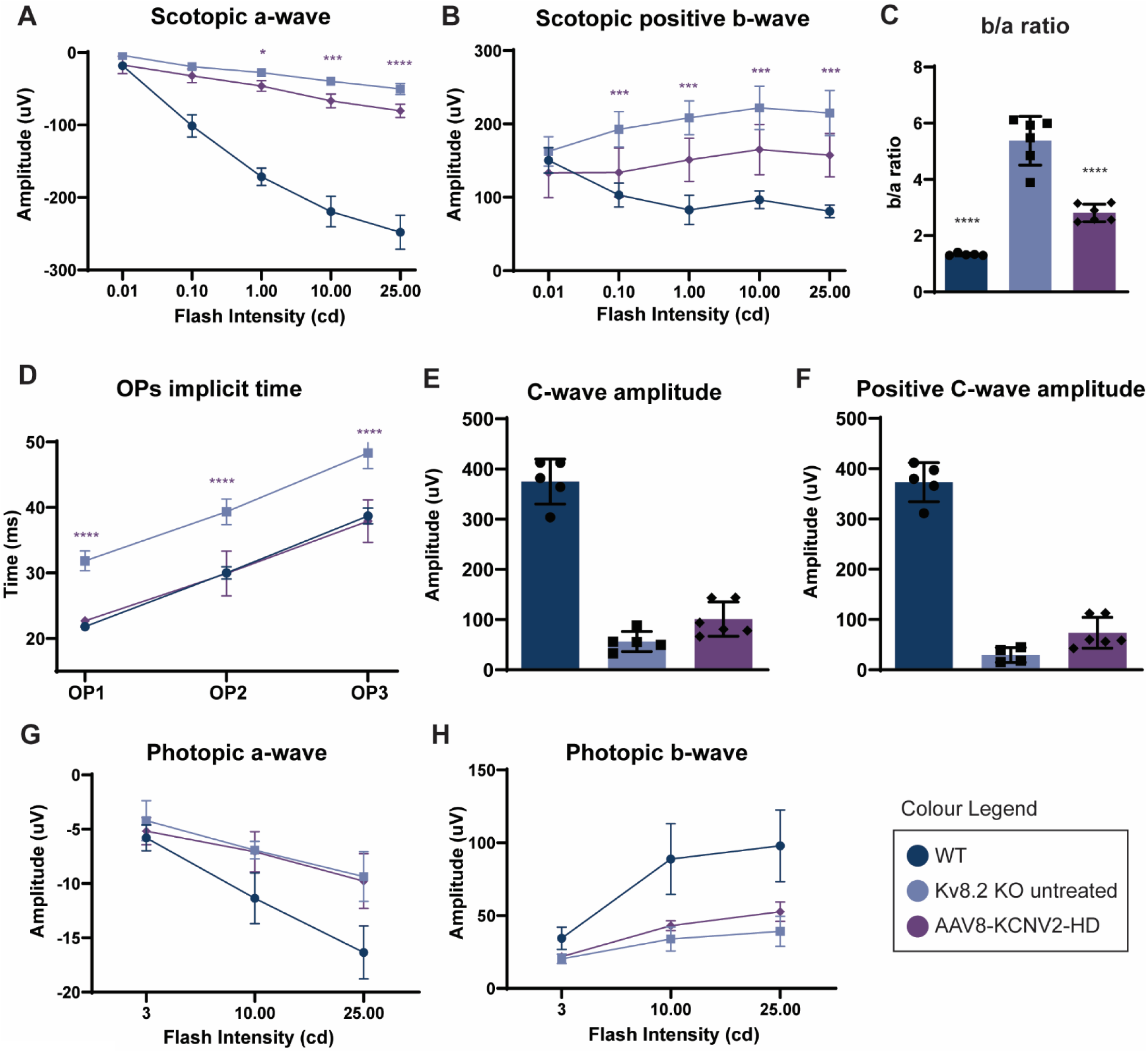
Long-term visual improvement is still present at 15 months post-treatment delivery. (A-B) Quantification at 15 months post-treatment of scotopic a-wave (A) and positive scotopic b-wave (B) at different intensities from animals treated at P30 with AAV8-KCNV2-HD and age-matched controls. **(C)** Quantification of scotopic b-wave to a-wave ratio at 25 cd.s/m^2^ alongside age-matched WT and untreated Kv8.2 KO mice. **(D)** Timing of oscillatory potential (OP) traces subtracted from the scotopic full flash responses at 25 cd.s/m2 for WT, Kv8.2 KO untreated and AAV8-KCNV2-HD treated mice. **(E-F)** Quantification of ERG c-wave responses showing amplitude measured from original baseline (E), amplitude measured from end of b-wave baseline (F). **(G-H)** Quantification of photopic a-wave (G) and b-wave (H) at different intensities. Numbers (n) are eyes from ≥ 3 biological replicates: WT n=5; Kv8.2 KO untreated n=6; AAV8-KCNV2 HD n=6. Graphs are averages ± SEM. Statistical analysis: two-way ANOVA followed by Tukey’s post hoc analysis. Significance is shown compared to Kv8.2 KO untreated mice: *p=0.0145; ***p=0.0002; ****p<0.0001.

### Treatment restores long term Kv8.2 subunit expression and correct localisation independent of age at treatment

Restoration of protein expression and its correct localisation after treatment is an essential step necessary to confirm correct protein translation from the delivered vector. In all our treated groups, expression of Kv8.2 was restored, and correct localisation confirmed at the inner segment of photoreceptors and its co-localisation with its co-partner subunit Kv2.1 (Fig 5A-C). Expression of Kv8.2 was restricted to the treated area (Fig 5B-C). These data show that an AAV-based gene replacement treatment is able to restore Kv8.2 expression, independent of age at treatment and time post-treatment. We were also able to confirm dose-dependent retinal expression of the codon-optimised human *KCNV2* gene delivered by our vectors (Fig 5D).

**Figure 5.**
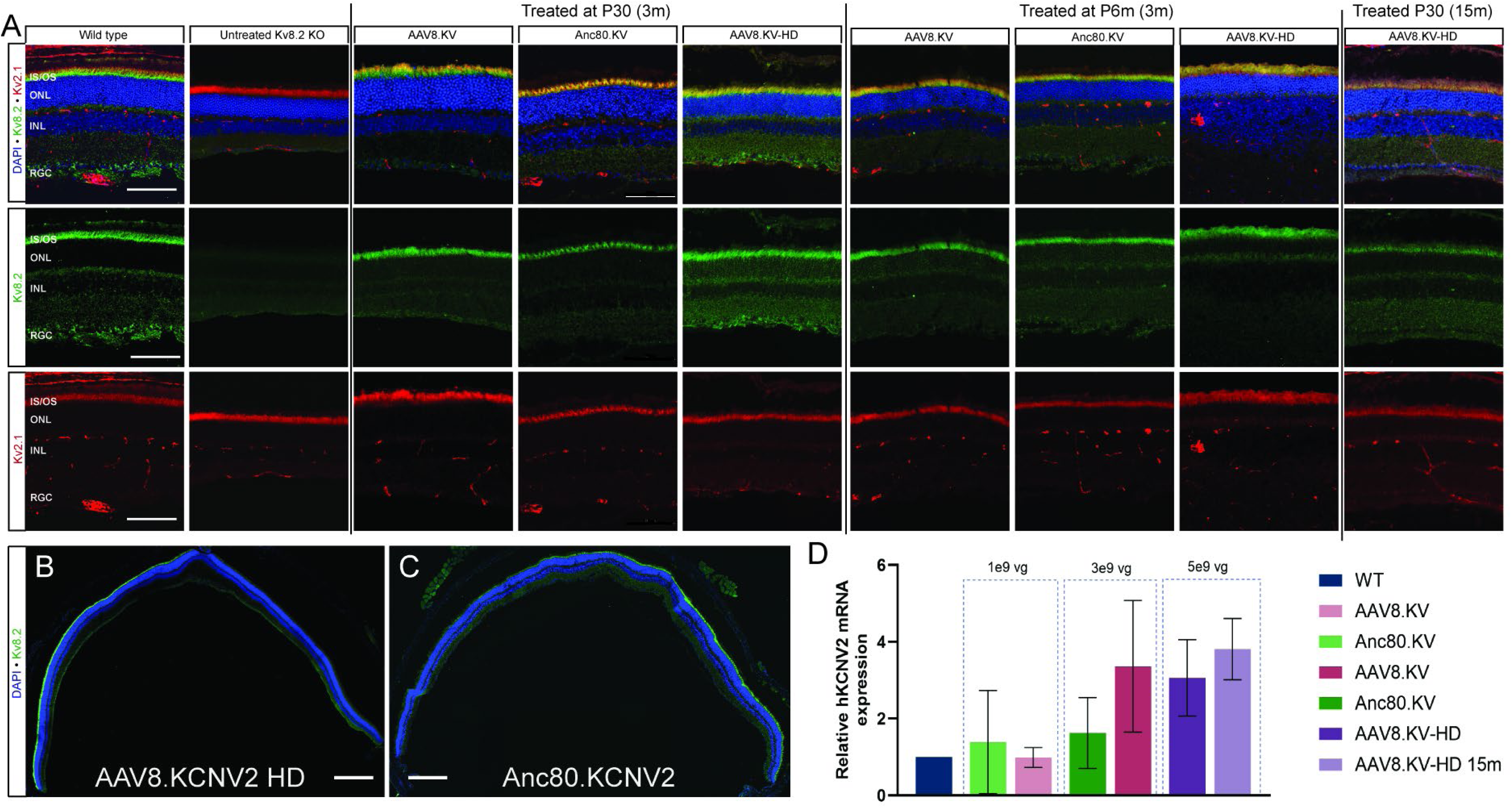
Retinal expression of Kv8.2 and co-partner Kv2.1 in the inner/outer segment of photoreceptors. **(A)** Representative images of retinal sections from eyes treated at different ages with AAV8.KCNV2, Anc80-KCNV2, and AAV8-KCNV2-HD, alongside WT and Kv8.2 untreated retinal sections labelled with Kv8.2 (green) and Kv2.1 (red) antibodies and DAPI (blue) nuclear stain. Treated samples are shown at age of treatment (P30 or P6m) followed by age of analysis (3m or 15m). Numbers of eyes analysed are from ≥ 3 biological replicates. Scale bar = 100µM. IS/OS, inner segment/outer segment; ONL, outer nuclear layer; INL, inner nuclear layer; RGC, retinal ganglion cell layer. **(B-C**) Tile images of example eyes treated with AAV8-KCNV2 HD or Anc80-KCNV2, showing spread of Kv8.2 expression (green) and cell nuclei layers (DAPI, blue). Scale bar = 300µM. **(D)** Real-time qPCR expression of codon-optimised human *KCNV2* gene inserted into different AAV vectors in treated eyes compared to mouse *Kcnv2* gene in WT mice. Graphs are averages ± SEM. Numbers (n) are eyes from 3-4 biological replicates for each group.

We have previously shown that Kv8.2 KO mice have an increased number of microglia cells in both their normal resting position in the inner nuclear layer, but also within the subretinal space where they become activated in response to the degeneration(28). As our treatment is viral based, it was important to check that the treatment was not increasing reactivity of retinal microglia. Figure 6 shows quantification of CD45+ cells (Fig 6A), overall microglia (Fig 6B) and both MCHII+ and CD11c+ microglia populations (Fig 6C-D) at both 1- and 3-months post-treatment, revealing no significant increase compared to untreated Kv8.2 KO. We also quantified histologically the amount of Iba1+ microglia in the inner retina and subretinal space; the treatment did not increase microglia numbers in either location (Fig 6E).

**Figure 6.**
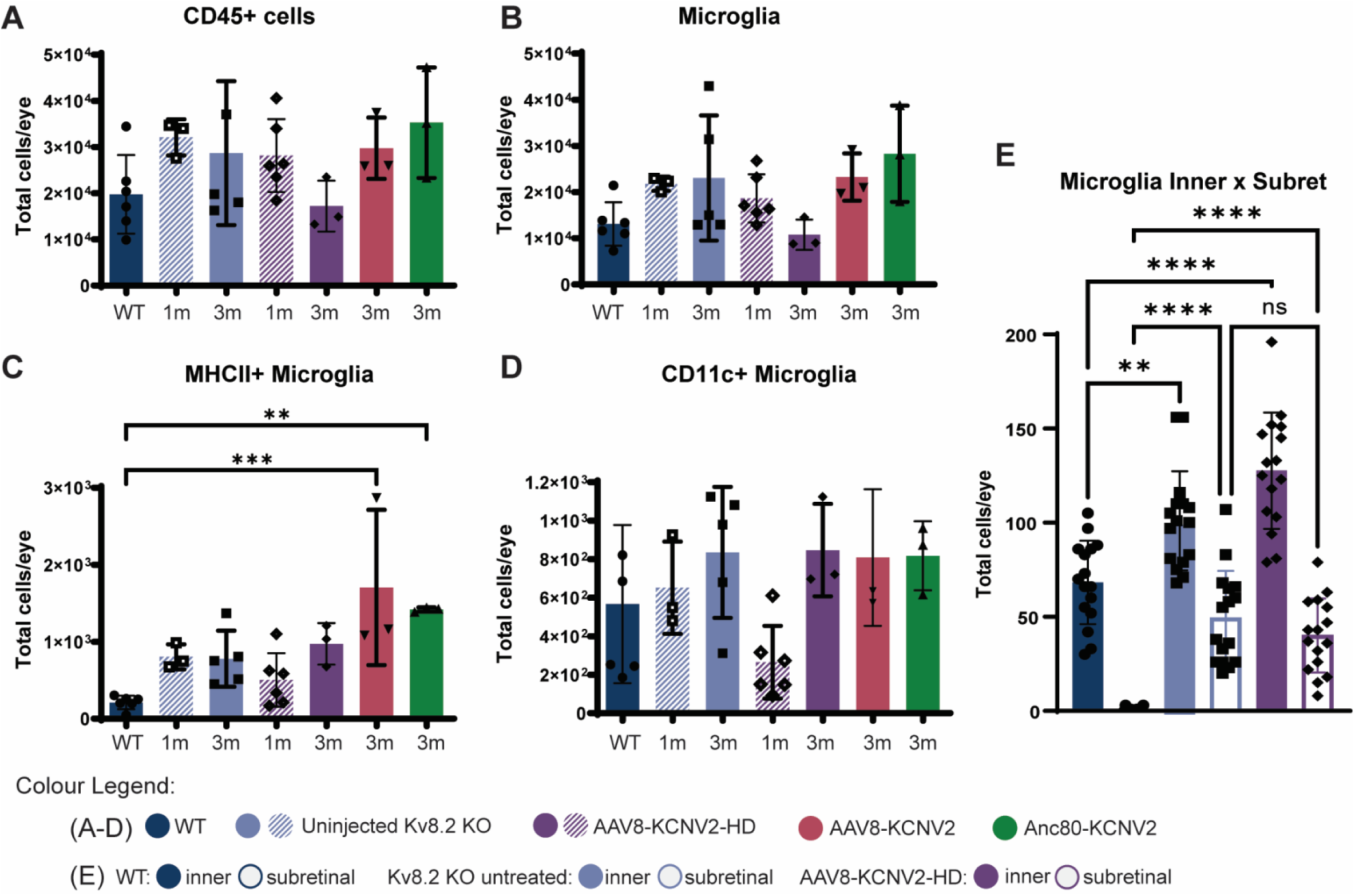
Treatment with AAV-KCNV2 vectors does not increase presence of immune cells in the retina. Graphs show cell numbers at 4 weeks (1m) and 12 weeks (3m) after treatment with AAV-KCNV2 vectors alongside age-matched WT and Kv8.2 KO untreated mice via flow cytometry (A-D) and histological quantification (E). **(A)** Total number of CD45-positive cells per eye. **(B)** Total number of Iba1+ microglia cells. **(C)** Total number of MCH class II positive microglia cells. **(D)** Total number of CD11+ microglia cells. Numbers (n) are per animal (2 eyes pooled): WT n=5-6; Kv8.2 KO untreated 1m n=3; Kv8.2 KO untreated 3m n=4-5; AAV8-KCNV2-HD 1m n=6; AAV8-KCNV2-HD 3m n=3; Anc80-KCNV2 3m n=3; AAV8-KCNV2 3m n=3. Bar graphs are averages ± SEM. Statistical analysis: one-way ANOVA followed by Tukey’s post hoc analysis. **p=0.0042; ***p=0.0004. **(E)** Histological quantification of number of Iba1+ microglia in the subretinal space and inner nuclear layer at 12 weeks post-treatment for the AAV8-KCNV2-HD groups compared to WT and Kv8.2 KO untreated retinas. N numbers indicate measurements from separate quadrants of the retina (4 quadrants/retina/area) from 4 retinas for WT, Kv8.2 untreated and AAV8-KCNV2-HD treated mice. Bar graphs are averages ± SEM. Statistical analysis: paired t-test **p=0.0034; ****p<0.0001.

### Single-cell RNASeq of Kv8.2 KO treated photoreceptors show specific targeting and gene expression changes in rod and cone photoreceptors

Single cell RNA sequencing was performed on whole retina dissociated cells from mice treated with AAV8.KV-HD and compared to age-matched untreated and wildtype controls. Cell clusters are displayed on a UMAP plot, with cell types identified by marker gene expression used in previous studies (56–58) (Fig 7A-B; Supplementary Figure 2). Bipolar cells were sub-grouped into rod, cone (OFF) and cone (ON) bipolar cells based on marker genes used by Shekhar et al. (2016)(59). The number of cells sequenced from each cell type is outlined in Suppl Table 1 and Suppl Figure 2D. From all analysed cells from the AAV8.KV-HD treated retinas, expression of human *KCNV2* was found predominantly in photoreceptors, with around 25% of both rods and cones expressing the transgene (Fig 7C). A small percentage of Muller glia (5%) and microglia (2.89%) also showed expression of the transgene, while all other retinal cells analysed had <2% expression (Fig 7C, Suppl Table 1).

**Figure 7.**
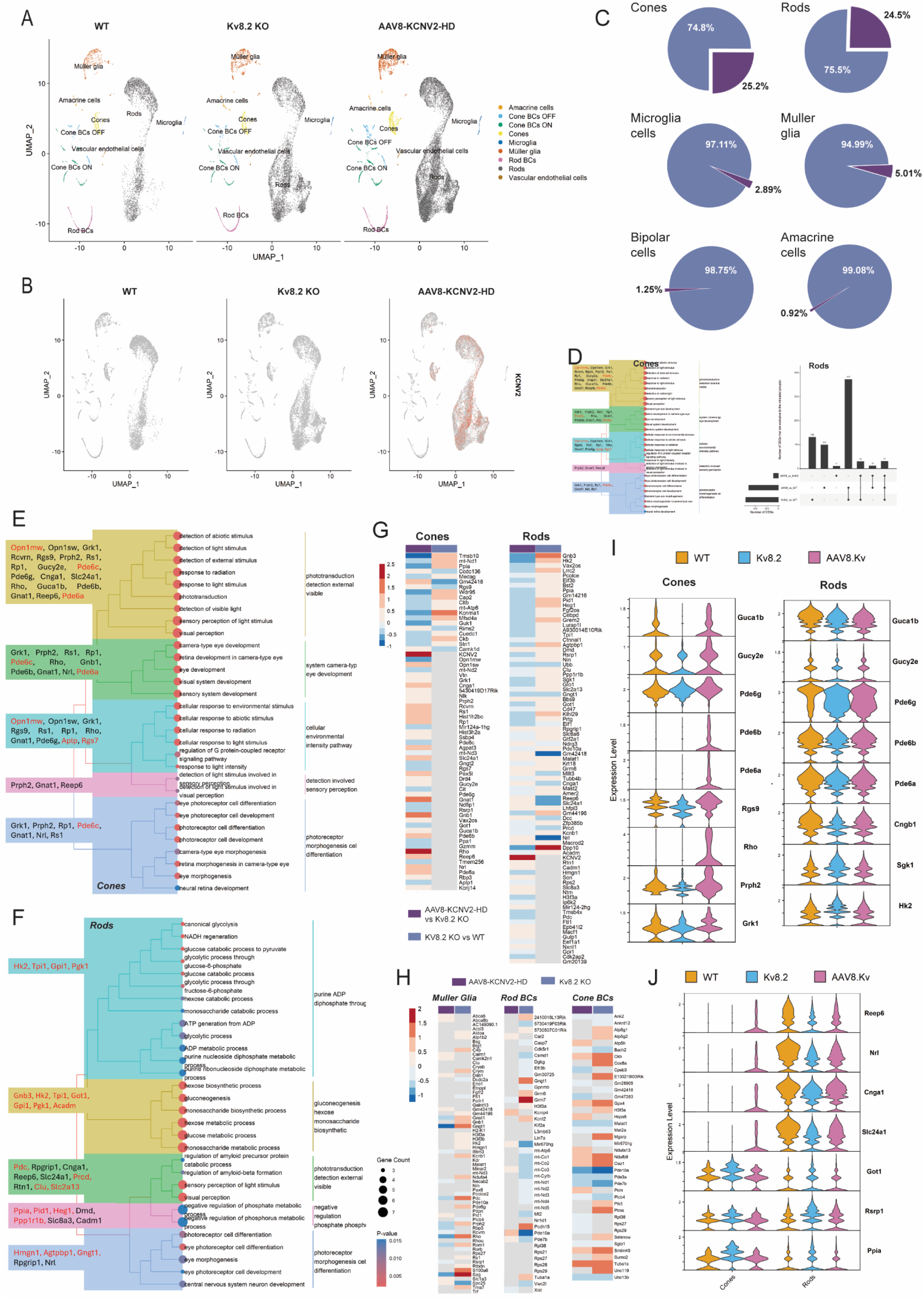
Single-cell RNA Sequencing reveals substantial changes in visual processing genes in rods and cones after treatment. **(A)** UMAP plot showing retinal cell types sequenced across the three groups. **(B)** UMAP plot of expression of codon-optimised human KCNV2 gene (co-hKCNV2+) across the three groups. Expression is restricted to the treated group AAV8-KCNV2-HD in rods and cones (red). **(C)** Pie charts of each retinal cell type showing percentage of co-hKCNV2+ cells within total of cells sequenced from AAV8-KCNV2-HD retinas. **(D)** Bar graphs indicating the number of differentially expressed genes (DEGs) in each group analysed in cones and rods. Lines indicate common DEGs across the linked groups. AAV8 indicates rods or cones from AAV8-KCNV2-HD retinas that were co-hKCNV2+. **(E)** Treeplot showing the top enriched GO biological processes associated with differentially expressed genes for co-hKCNV2+ cones from the AAV8-KCNV2-HD treated retinas compared to untreated Kv8.2 KO cones. **(F)** Treeplot showing the top enriched GO biological processes associated with differentially expressed genes for co-hKCNV2+ rods from the AAV8-KCNV2-HD treated retinas compared to untreated Kv8.2 KO rods. **(G)** Heatmap comparison of cones and rods DEGs from AAV8-KCNV2-HD compared to untreated Kv8.2 KO, and Kv8.2 KO compared to WT. **(H)** Heatmap of DEGs from Müller glia, rod bipolar cells (BCs) and cone BCs comparing between these cell types in AAV8-KCNV2-HD treated retinas to Kv8.2 KO untreated. **(I)** Violin plots of specific DEGs in cones and rods from AAV8-KCNV2-HD (co-hKCNV2+ only), WT and Kv8.2 KO untreated retinas. **(J)** Violin plots of DEGs that are common to both co-hKCNV2+ cones and rods from AAV8-KCNV2-HD retinas compared to expression of same genes in WT and Kv8.2 KO untreated retinas.

Rods and cones expressing the human *KCNV2* transgene showed 84 and 65 differentially expressed genes (DEGs) compared to untreated, respectively (Fig 7D). The top enriched gene ontology (GO) biological processes for treated cones compared to untreated are related to visual perception and light stimulus, including phototransduction and are shown in Fig 7E alongside the genes linked to these top biological processes. Of the 65 DEGs in cones (Fig 7G), 36 genes have upregulated expression compared to untreated Kv8.2 KO cones, with 31 involved in either the visual cycle, retinal development, or known to cause IRDs if mutated. We also saw an upregulation of rod isoforms of phototransduction genes in the treated cones (Fig 7G), including *Rho, Gnat1, Cnga1, Pde6a* and *Pde6b*. However, cross-contamination from rods is a known issue in single-cell retinal samples(60). We also saw a significant upregulation of s-opsin (*Opn1sw*, Fold Change [FC] 1.45; *p*<0.0001), but a downregulation of m-opsin (*Opn1mw*, FC 0.70, *p*<0.0001) in cones after treatment. Apart from *KCNV2* upregulation, we also noticed changes in treated cones in genes involved in potassium ion transport: calcium-activated potassium channel subunit alpha-1 (*Kcnma1,* FC 0.66; *p*<0.0001), potassium-dependent sodium/calcium exchanger (*Slc24a1,* FC 1.68; *p*=0.0004), regulator of G-protein signalling 7 (*Rgs7,* FC 0.76; *p*=0.0007) and the potassium inwardly-rectifying channel, subfamily J, member 14 (*Kcnj14,* FC 1.22; *p*=0.012). From the 29 downregulated genes, 11 are involved with visual function or development, while around 9 are involved in mitochondria function/energy homeostasis.

In treated rods, the top enriched GO biological processes compared to untreated rods were related to phototransduction and visual detection, ATP biosynthesis and aerobic respiration (Fig 7F). From the 84 DEGs (Fig 7D), 27 were upregulated, while 57 were downregulated after treatment. From the total DEGs in rods, 39 have confirmed action, expression or involvement with retinal function, degeneration or development. Interestingly, treated rods showed a significant downregulation of the *Kcnb1* gene which encodes for Kv8.2 co-partner Kv2.1 (FC 0.87, *p*=0.0057; Fig 7G). This is in equal opposition to the upregulation of *Kcnb1* expression in untreated Kv8.2 KO rods compared to WT (FC 1.16; *p*<0.0001). This could be explained by the downregulation of *Sgk1* (serum/glucocorticoid regulated kinase 1) after treatment (FC 0.78, *p*<0.0001; Fig 7I), and its upregulation in untreated Kv8.2 KO rods compared to WT (FC 1.33; *p*<0.0001). *Sgk1* is known to be upregulated during cellular stress and energy depletion, and to directly regulate the expression of Kv subunits(61). Furthermore, the treatment was also able to downregulate the expression of *Hk2* and *Got1* genes in rods (Fig 7I-J). The Hk2 enzyme (Hexokinase 2, FC 0.65; *p*<0.0001; untreated Kv8.2 x WT FC 1.54; *p*<0.0001), is one of the main regulators of aerobic glycolysis, and apoptotic signalling(62), while *Got1* (Glutamic-oxaloacetic transaminase 1, FC 0.83; *p*<0.0001; untreated Kv8.2 x WT FC 1.20; *p*<0.0001) is involved in glycolysis pathways.

A small number of up- and down-regulated genes were common to treated rods and cones (Fig 7J). These include upregulation of *Reep6, Nrl, Slc24a1* and *Cnga1*, and downregulation of *Got1, Rsrp1* and *Ppia*. In adult rods, *Nrl* (Neural leucine zipper) expression is essential for homeostasis, and its expression is downregulated in untreated Kv8.2 KO rods (FC 0.57; *p*<0.0001) and absent in untreated cones. Interestingly, Nrl is known to drive the expression of *Reep6* (Receptor Accessory Protein 6), responsible for protein trafficking from the ER to rod OS(63). Loss of *Reep6* leads to retinal degeneration and disruption of ER homeostasis and impairment of guanylate cyclase trafficking to the OS, and thus it has an important role in maintaining cGMP homeostasis(63). Indeed, in untreated Kv8.2 KO rods, both *Reep6* and *Guca1b* (Guanylate Cyclase Activator 1B) are downregulated (FC 0.63 and FC 0.80, respectively; *p*<0.0001). In cones, although *Reep6* expression is not significantly altered in untreated Kv8.2 KO, it is upregulated after treatment, most likely due to increased *Nrl* expression, and correlates to the upregulation of *Guca1b* (FC 1.37; *p*=0.0092)*, Gucy2e* (Retinal guanylyl cyclase 1, FC 1.25; *p*=0.00035), rod phosphodiesterase 6 subunits a (*Pde6a*, FC 1.60; *p*=0.036), b (*Pde6b*, FC 1.46; *p*=0.0095) and g, (*Pde6g*, FC 1.24; *p*=0.00040), and *Cnga1* (Cyclic nucleotide gated channel subunit alpha 1, FC 1.56; *p*=0.00042), which are all localised in the OS.

Treatment effects on transcriptomic changes were not restricted just to the targeted photoreceptor cells. We also noticed a shift in overall gene expression in Müller glia, and in cone and rod bipolar cells, where after treatments, genes that were significantly affected in untreated Kv8.2 KO cells showed a shift in the opposite direction of expression (Fig7H). Interestingly, and similar to a previous study(64), we observed expression of several visual function and phototransduction genes in Müller glia cells. However, when comparing the number of Müller glia DEGs in untreated Kv8.2 KO vs WT to treated Kv8.2 KO to WT, there was a significant reduction in the number of expressed genes (303 vs 59), potentially indicating less reactive gliosis after treatment (Fig 7G). Furthermore, the compensatory expression of photoreceptor-related genes in Müller glia appears to be corrected after treatment (Fig 7G).

### Whole retina proteomics shows increased cell function, metabolism and survival

Whole retinal tissue proteomics was performed in wildtype (n=3), untreated Kv8.2 KO (n=3) and AAV8.KV-HD treated Kv8.2 KO mice (n=4) and comparison between treated and untreated retinas revealed significant changes (>1.1-fold, *p*<0.05) in 124 proteins, evenly split with 62 proteins upregulated and 62 downregulated (Fig 8A-C). The Principal Component Analysis (PCA) plot shows a clear separation between the treated and untreated samples along the principal component 1 (PC1), which explains more than 70% of the total variance, indicating that this component strongly differentiates the two groups. Within each sample type, the replicates were clustered together, suggesting good reproducibility within the groups. Identification of differentially regulated proteins (DEPs) using a volcano plot (Fig 8B) and heatmap (Fig 8C) also reveal significant changes in protein expression between the treated and untreated samples. From the protein-protein interaction network, it is clear that several metabolic and protein biosynthesis and complex binding are altered after treatment (Fig 8D). The top 10 enriched GO terms are shown for Biological Processes (BP; Fig 8E), Cellular Component (CC; Fig 8F), Molecular Function (MF; Fig 8G) and Reactome Pathways (Fig 8H).

**Figure 8.**
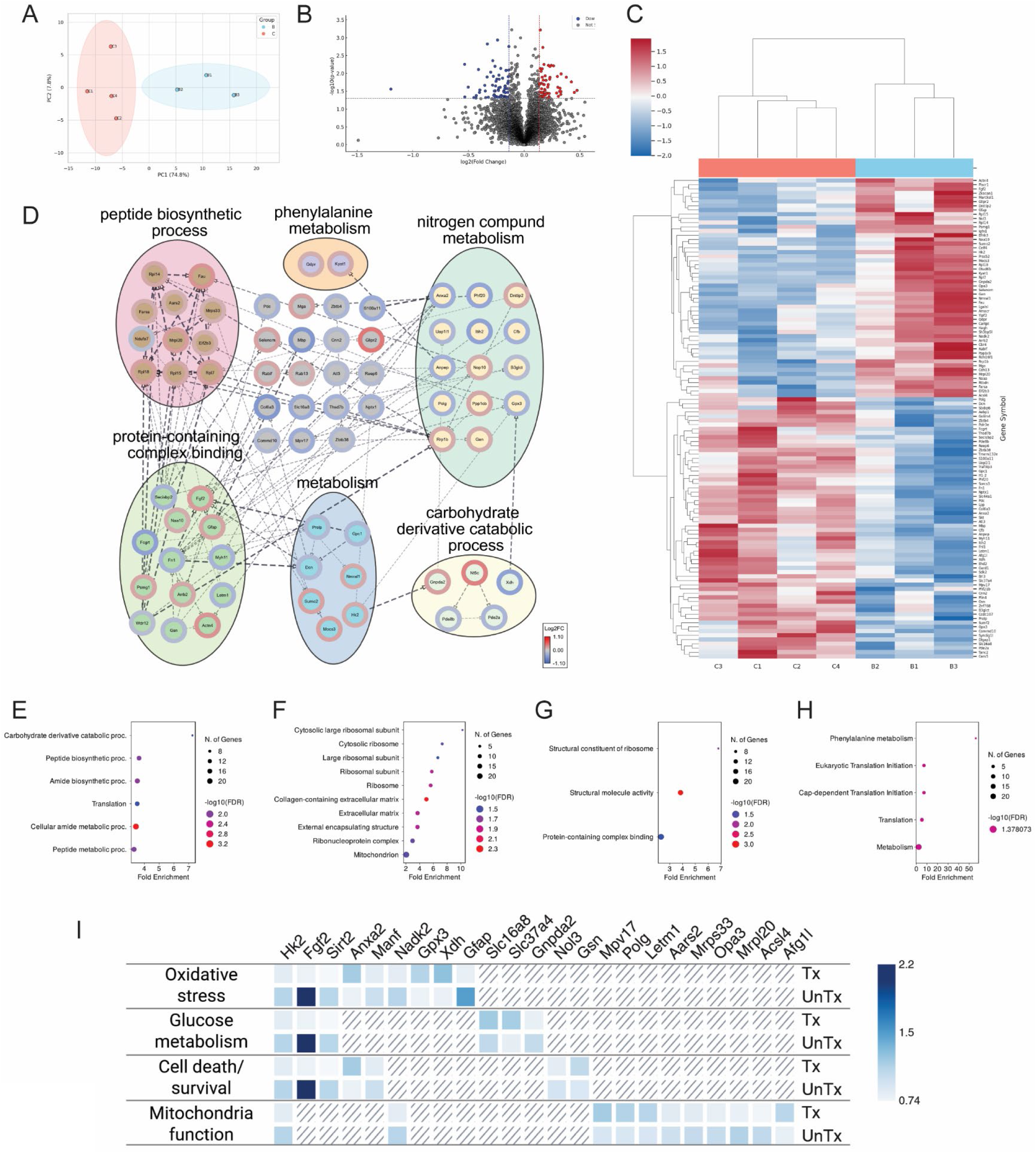
Whole retina proteomics reveals improved metabolic changes after treatment. **(A)** PCA plot showing a clear separation between the two sample types (B, Kv8.2 KO untreated retinas (n=3) and C; AAV8-KCNV2-HD treated retinas (n=4)) along the principal component 1 (PC1). **(B)** Volcano plot of differentially regulates proteins (DEPs) of AAV8-KCNV2-HD treated retinas compared to Kv8.2 KO untreated retinas. Each point corresponds to a quantified protein, with the −log₁₀(p-value) on the y-axis and the fold change (b/c) on the x-axis. Vertical dashed lines define the fold change threshold of 10%. Proteins exceeding both the significance threshold and fold change are considered DEPs, with down-regulated proteins shown in blue and up-regulated proteins in red. (C) Heatmap displaying the relative abundance of differentially expressed proteins (DEPs) in AAV8-KCNV2-HD treated retinas (red bar, n=4) compared to Kv8.2 KO untreated retinas (blue bar, n=3). Rows represent specific proteins, and columns represent different samples. The color scale represents the relative abundance of detected proteins in each sample, ranging from blue to red indicating higher abundance. The hierarchical clustering was performed on both rows and columns and are shown as dendrograms on the left and top, respectively. **(D)** Protein-protein interaction network constructed using the stringApp plugin in Cytoscape for 124 differentially expressed proteins (DEPs) represented by their gene symbols. Only medium-confidence clusters (STRING score > 0.4) containing at least two nodes are displayed. The stringApp plugin identified significantly enriched (FDR < 0.05) Biological Processes (BP), Molecular Functions (MF), and Reactome Pathways for clusters, which were then color-coded as pie chart illustrating the fold changes of the DEPs using the OmicsVisualizer plugin. Dashed edges indicate experimentally derived interactions between DEPs. (E-H) Top 10 enriched GO terms in AAV8-KCNV2-HD treated retinas compared to Kv8.2 KO untreated retinas. Terms in Biological Processes (E), Cellular Component (F), Molecular Functions (G) and Reactome Pathway (H) were sorted by the fold enrichment on the x-axis. The size of the circle represents the quantity of mapped proteins involved in each GO. Red colour indicates increasing -log10 (FDR) and blue colour shows decrease. A total of 124 protein IDs were mapped to Mus Musculus genes and were analyzed using ShinyGO 0.82. The significant GO terms with an FDR less than 0.05 and without redundancy were presented here. **(I)** Overview of common differentially expressed genes (DEGs) and DEPs in AAV8-KCNV2-HD treated compared to Kv8.2 KO untreated samples showing selected genes/proteins and their associated biological function.

As these results are not cell-specific but instead reveal changes in the retina as a whole, including treated and untreated photoreceptors, we were not able to confirm most of the photoreceptor-specific changes we saw in the scRNASeq data. One significant similarity between the datasets was the upregulation of REEP6 and downregulation of HK2 and FGF2 after treatment. Interestingly, phosducin (PDC) was shown to be upregulated in our proteomics dataset (Fig 8I), but in our single-cell RNASeq data it was upregulated in cones, but downregulated in rods, Muller glia and amacrine cells after treatment. Phosducin is a modulator of the phostotransduction cascade – by binding to the beta-gamma subunits of the retinal G-protein transducin in photoreceptors, it helps to keep the alpha subunit active for longer (65).

Further analysis of the differentially expressed proteins in AAV8.KV-HD retinas compared to untreated showed significant differences in proteins involved in oxidative stress, mitochondria function, glucose metabolism and cell death/survival pathways. When compared to Kv8.2 KO untreated versus wildtype expression data, the changes in expression of almost all of these selected proteins are reversed after treatment (Fig 8I). After treatment, there was a downregulation of GFAP (glial fibrillary acidic protein), FGF2 (fibroblast growth factor 2), SIRT2 (NAD-dependent protein deacetylase sirtuin-2), HADK2 (NAD kinase 2), NOL3 (nucleolar protein 3) and MANF (mesencephalic astrocyte-derived neurotrophic factor), all proteins involved in protecting against cellular stress and oxidative damage, indicating a potential recovery of cells towards homeostasis. Interestingly, our data showed both upregulation and downregulation of proteins involved in glucose metabolism, essential for photoreceptors function. Both Slc16a8 and Slc37a4, members of the solute carrier family 16 member 8, and family 37 member 4, respectively, were upregulated in treated retinas. Slc16a8, also known as MCT3 (Monocarboxylate Transporter 3), is involved in the transport of lactate and other monocarboxylates and important for metabolic coupling between retinal pigment epithelium and photoreceptors. Slc37a4, also known as G6PT1 (Glucose-6-Phosphate Transporter 1), is involved in glucose-6-phosphate transport and is critical for maintaining glucose homeostasis. However, our data also showed a decrease in the expression of HK2 (hexokinase 2), an activator of aerobic glycolysis, and GNPDA2 (Glucosamine-6-Phosphate Deaminase 2), part of the hexosamine biosynthetic pathway. This discrepancy in glucose metabolism could be explained by proteomics reflecting the overall averaged expression that included treated and untreated cells. It is also seen in mitochondria function related proteins, where protein synthesis is downregulated (MRPL20, MRPS33, AARS2), versus DNA replication and repair (MPV17, POLG), which were upregulated.

Another point of interest was the significant changes in expression of SIRT2 and NADK2 post-treatment. They are both upregulated in untreated retinas and downregulated after treatment (Fig 8I). SIRT2, or Sirtuin 2, is an NAD^+^ (nicotinamide adenine dinucleotide)-dependent deacetylase, while NADK2 or NAD kinase 2, is a mitochondrial enzyme that phosphorylates NAD^+^ to NADP^+^, fuelling cellular energy. Given the dependency of SIRT2 on NAD+ for function, it was interesting to note in our data a potential correlation of their expression levels.

### Treatment restores Kv8.2 protein in patient-derived retinal organoids

In order to assess whether our treatment was also capable of driving Kv8.2 expression and restoring Kv8.2 protein in a human-based model, we delivered our treatment to retinal organoids derived from a patient iPSC line carrying the pathogenic *KCNV2* variants c.[775G>A][782C>A] p.[Ala259Thr][Ala26I Asp]. At 26 weeks, *KCNV2^775G>A/782C>A^* patient-derived retinal organoids developed similarly to healthy control retinal organoids, showing the presence of both cone photoreceptor cells (ARRESTIN3, ARR3+ cells) and RHODOPSIN+ rod photoreceptor cells. Furthermore, organoids developed robust outer segments (PERIPHERIN+)(Fig 9A-C). Analysis for *KCNV2* mRNA showed that both control and *KCNV2^775G>A/782C>A^* organoids expressed similar levels of *KCNV2* (Fig 9D, n= 20 organoids, N= 2-4 differentiation batches), given that the variants this patient harbours are missense mutations.

**Figure 9.**
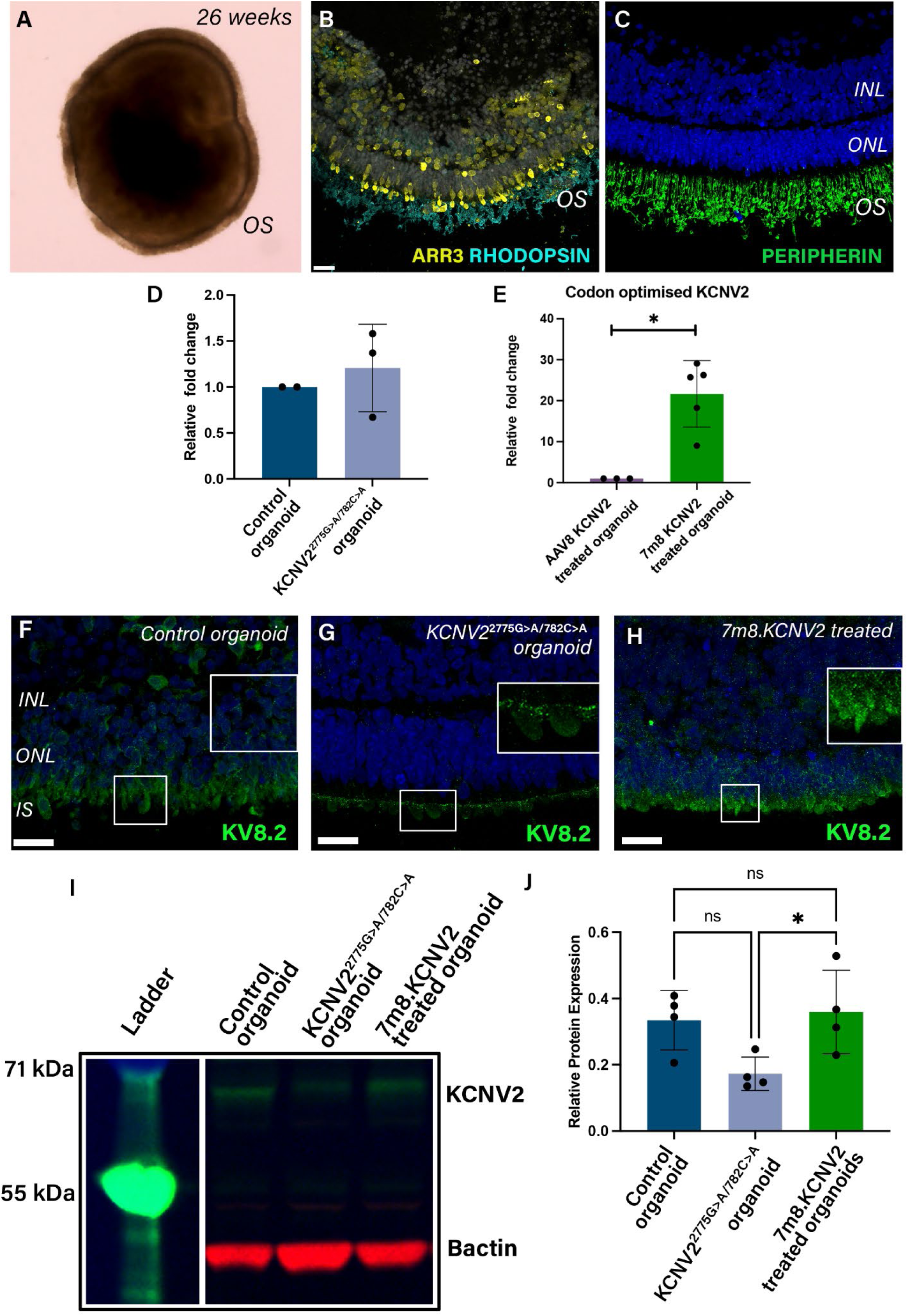
AAV-based Gene therapy efficiently transduces *KCNV2* patient-derived retinal organoids. (A-C) Representative images of retinal organoids (RO) derived from a patient harbouring the 775G>A;782C>A disease variants in *KCNV2*. (A) Phase contrast image of a 26 weeks old KCNV2 patient RO showing the outer segment (OS) brush border. (B-C) histological section image of a KCNV2 patient RO labelled with photoreceptor markers cone Arrestin (ARR3, yellow), Rhodopsin (cerulean) and Peripherin (green). Cell nuclei are shown in blue (DAPI). **(D-E)** Real-time qPCR measuring mRNA expression of native human KCNV2 transcript in WT and untreated ROs (D), and codon-optimised human KCNV2 transcript in AAV8 or AAV7m8 treated ROs (E). Graphs are averages ± SD. Statistical analysis: Mann-Whitney t-test \**p*=0.0357. **(F-H)** Representative histological images of ROs showing expression of Kv8.2 (green) in the inner segment (IS) region of WT, untreated KCNV2, and AAV7m8-KCNV2 treated ROs. Scale bar = 20µm. **(I-J)** Western blot quantification of Kv8.2 expression from WT, untreated and treated ROs. The number of organoids analysed are from ≥ 3 ROs differentiation replicates. Graphs are averages ± SD. Statistical analysis: one-way ANOVA followed by Tukey’s post hoc analysis: \**p*=0.049.

Next, we treated *KCNV2^775G>A/782C>A^* organoids at 20 weeks with either AAV8 or AAV7m8 vectors and samples were collected for analysis 6 weeks post-treatment (26 weeks). Given that AAV transduction of retinal organoids is best achieved with the AAV7m8 serotype(66), it was not surprising that organoids treated with the AAV8.KCNV2 vector showed little to no expression of the human codon-optimised *KCNV2* transgene, while organoids treated with AAV7m8 showed a 2.5-fold increase in expression compared to control organoids (Fig 9E, n= 15 organoids, N= 3-5 differentiation batches).

Next, we performed immunostaining for KV8.2 showing its punctuated expression in the inner segment region of photoreceptor cells. Interestingly, in the *KCNV2^775G>A/782C>A^* protein localisation appeared ectopically localised, while treatment with 7m8.KCNV2 led to a more widespread expression of the protein (Fig 9F-H). Finally, western blot quantification showed a significant increase in expression of Kv8.2 protein in treated organoids with protein levels similar to control levels (Fig 9I-J, n= 20 organoids, N= 4 differentiation batches). These data demonstrate that our RK.*KCNV2* viral vector construct effectively restores KV8.2 protein expression in human photoreceptor cells.

## Discussion

CDSRR or *KCNV2* retinopathy is part of the incurable group of conditions referred to as inherited retinal diseases (IRDs). It is caused by mutations in the *KCNV2* gene that directly impact the function and survival of the light-sensing photoreceptor cells, specialised neurons that do not regenerate. CDSRR patients suffer from a progressive loss of photoreceptors starting in early childhood, and this loss will eventually lead to irreversible vision loss during adulthood and legal blindness later in life. There is no treatment available or any prophylactic management options to slow loss of vision. Furthermore, *KCNV2* retinopathy is also clinically classified as a cone-rod dystrophy, which means that the cone-driven visual system is affected early in the disease. The cone-driven visual response is what enables focus, perception of colours and bright lights, and is responsible for much of our everyday tasks such as reading, driving and recognising faces. The loss of cone-mediated vision is, therefore, extremely detrimental to patients, having a high impact on their quality of life and independence.

Developing effective treatments for IRDs has significant challenges due to its clinical and genetic heterogeneity, and the orphan disease status of each condition. With mutations in over 300 genes are known to cause IRD (retnet.org), progress has been slow and understandingly frustrating to patients. However, due to the monogenic nature of IRDs, accessibility and size of the eye, the application of viral vector-based gene therapy treatments has seen unprecedented successes in this area, culminating in the first FDA-approved *in vivo* gene therapy in 2017 (Luxturna®) for IRD patients with mutations in the *RPE65* gene. Other treatment approaches being investigated for IRDs include neuroprotection therapies, which aim to preserve retinal cells and could benefit a wider range of patients but are limited in their capacity to restore full cellular function or treat the underlying cause. The advantage of gene therapy lies in its potential to cure the condition by replacing the defective gene. In this study, we demonstrated for the first time that a gene replacement therapy for *KCNV2* retinopathy can replace and restore the function of the defective Kv channels in photoreceptors, leading to significant improvements in visual function. Our treatment was able to successfully and exclusively target cone and rod photoreceptors – the only retinal cells that express *KCNV2* (33, 34). The successful targeting of our treatment was confirmed not only by changes in cellular activity via ERG responses, but also via transcriptional changes in RNA and protein levels. The selection of the human rhodopsin kinase promoter was crucial in ensuring correct targeting of our treatment exclusively to rods and cones. This promoter has been extensively used in preclinical gene therapy studies to restrict expression to photoreceptors (67–71), and is also the promoter of choice in current gene therapy clinical trials for IRDs, further validating the clinical translatability of our treatment.

Expression of the Kv8.2 subunit has been shown to specifically co-localise to the inner segment of the photoreceptors and co-expressing with Kv2.1 on the segment membranes (28, 33, 34). In the Kv8.2 KO mice, expression of Kv8.2 is completely absent while Kv2.1 is still present in the inner segments. This is not unexpected, as Kv2.1 can form homomeric channels on their own, while Kv8.2, being a silent subunit, cannot traffic to the membrane without a Kv2 subunit present (34). This explains why the Kv2.1 KO mice lack both Kv2.1 and Kv8.2 expression (28, 33). The lack of the modulatory Kv8.2 subunit is also thought to be behind the dysregulated electrophysiological response seen in the Kv8.2 KO mice and *KCNV2* patients. The Kv channels are responsible for setting the dark resting current in photoreceptors, which in conjunction with the Na^+^/K^+^ exchanger, maintains a steady flux of K^+^ ions in the dark. Light activation of photoreceptors causes the membrane potential to change, and this change is sensed by the ion conducting and voltage sensor core, causing the Kv channel to close. The activation and deactivation dynamics of heteromeric Kv2.1/Kv8.2 and homomeric Kv2.1 channels are known to differ significantly (24), which explains why visual response is not completely abolished in the absence of Kv8.2 subunits, as homomeric Kv2.1 channels might still be responsive to light activation. Indeed, the timing of a- and b-wave peaks are delayed in *KCNV2* patients (1, 2, 29), in line with a slower activation response of homomeric Kv2.1 channels to membrane potential changes (24). The potential role of homomeric Kv2.1 channels on pathophysiology of *KCNV2* retinopathy remains unclear. However, our data show that *Kcnb1* expression is significantly increased in Kv8.2 KO rods compared to WT, while AAV8 treated rods show a significant decrease of *Kcnb1* mRNA compared to untreated, indicating a restoration of hom/het Kv2.1/8.2 channels balance in the inner segments of treated rod photoreceptors.

These results clearly demonstrate that our gene therapy treatment significantly improved the response of both rods and cones to light and restored vision to the treated KO mice. Vision restoration was seen on all levels of the visual transduction pathway, from cellular (restored Kv8.2 expression in photoreceptor’s inner segments) to localised retinal function (via ERG responses) to higher visual pathways via restoration of the optomotor response. The full-field ERG response presented in this study is an averaged response from the different retinal cells across the whole retina. This inevitably shows a reduced response compared to wildtype levels as the treatment area is restricted – as is in the case of subretinal injection. This masks the treatment efficacy within its treated area, since area size is known to correlate to ERG amplitude (72). Thus, using different measures of visual response that go beyond retinal activity can provide a more comprehensive understanding of treatment effectiveness and impact on visual tasks (73). The results show that our treatment was capable of fully recovering the optomotor response of treated animals to wildtype levels. This outcome is maintained even when treatment is administered much later in the disease progression, suggesting that treatment of even a small number of photoreceptors can provide restoration up to normal vision levels.

Some improvements in vision can still be seen even when treatment is administered at later stages of disease, most likely due to the slow rate of photoreceptor degeneration present in this condition (28, 33). However, to achieve significant outcomes for all measures, particularly the different ERG responses, earlier treatment administration appears to be more effective. The effectiveness of early treatment was recently showed to be crucial in restoring activation of the visual cortex in achromat patients that received a subretinal AAV gene therapy (74, 75). Different from *KCNV2* retinopathy, achromatopsia is characterised by the absence of colour vision and acuity from birth, which raises the question whether higher cortical responses can be modulated after ocular gene therapy. Indeed, preclinical studies showed that optomotor response recovery post-gene therapy in a mouse model of achromatopsia was age dependent (76). This would, however, not pose too much of a limitation on *KCNV2* patients, as the slow progression of the disease likely indicates that higher cortical pathways are still preserved (1, 2). Our data appears to be in line with this, as restoration of normal vision via optomotor reflex testing was seen in all age treated groups. This could indicate that there is sufficient preservation of the cortical pathways even at late stages of disease, and that treatment of even a small number of photoreceptors could provide recovery to a perceived normal level of visual response in patients. However, evaluation of visual cortex activity in *KCNV2* patients has not been carried out, so it remains unclear how much modulation occurs during disease progression, and if there is an age effect that could impact treatment outcomes. Nevertheless, our data indicate that the effectiveness of treatment is age-dependent, but even when treatment is administered later in the disease stage, improvements in visual function are still possible.

Beyond the effective recovery of visual function, our data show that the treatment is safe. Our results showed no detrimental effects of the gene therapy, either via *in vivo* functional assessments, transcriptional analysis or when tested in patient-derived retinal organoids. Subretinal AAV gene therapy is generally considered safe and tolerable, with trials demonstrating positive safety and efficacy outcomes (40). However, potential risks and complications exist, including immune responses, targeting errors, and the possibility of causing unintended long-term effects (77–79). Luxturna®, the first AAV gene therapy approved by the FDA, has now been commercially available in the USA since 2017, with patients from the first clinical trials having received the therapy in 2007(80–82). Despite the known risks and safety concerns, gene replacement therapy for several types of IRDs is the only potential curative treatment approach. This study and several others, provide strong evidence that visual rescue can occur after AAV-based gene therapy treatment.

In summary, in this study we demonstrate that gene replacement therapy can either fully or partially correct all features of this disorder mirrored in the Kv8.2 KO mouse model. Given the relatively slow progression of this condition, the presence of a high number of photoreceptors even in older patients and its distinctive diagnostic ERG profile (1, 2, 29), *KCNV2* retinopathy presents as an ideal candidate for future clinical trials to test the efficacy of this type of curative treatment approach. The relevance of this study is further enhanced by comparative data from both *in vivo* mouse and human-based disease models, a first step in addressing the high failure rate of drugs in clinical trials due to efficacy concerns (83). By testing our therapeutic approach in diseased human retinal organoids, we provide comprehensive pre-clinical efficacy data across species. This is also in alignment with the FDA’s recent directive on organoid toxicity testing (New Approach Methodologies or NAMs data). This dual-model strategy significantly enhances the robustness and translational potential of the research findings generated by this study, paving the way for a future clinical trial for *KCNV2* retinopathy.

## Materials and Methods

### Vector generation and production

The DNA insert containing the short rhodopsin kinase promoter (RK)(39) followed by either a codon-optimised version of the human *KCNV2* gene (Ensembl transcript ENST00000382082.4 KCNV2-201) or the enhanced green fluorescent protein (eGFP) gene, a WPRE sequence (84) downstream of either gene, and a BGH polyA was synthesised by GeneWiz (South Plainfield, USA) and cloned into a pAAV plasmid backbone. The resulting pAAV-RK-hKCNV2 or pAAV-RK-eGFP plasmids were used to produce Anc80L65.RK.hKCNV2 (referred as Anc80-KCNV2), AAV8.RK.hKCNV2 (referred as AAV8-KCNV2), Anc80L65.RK.eGFP (referred as Anc80-GFP) and AAV8.RK.eGFP (referred as AAV8-GFP) viral vectors at the Gene Transfer Vector Core facility (Grousbeck Gene Therapy Center, Ocular Genomics Institute of Harvard Medical School, and Massachusetts Eye and Ear Infirmary, Boston, USA). Final titres were 3x10^12^ and 5x10^12^ vector genomes (vg)/ml for Anc80-KCNV2 and AAV8-KCNV2, respectively. Appropriate dilutions were made in separate aliquots on the day of injection for different titres tested. iPSC-derived retinal organoids (ROs): AAV8-KCNV2 and AAV7m8-KCNV2 vectors used for transduction of ROs was made by Vector Building using the same transgene construct described above. Final titres were 2.8x10^13^ vg/ml (AAV8) and 2.9x10^13^ vg/ml (AAV7m8).

### Subretinal injections

Mice were anesthetised with an intraperitoneal injection of 80mg/kg Ketamine (Ceva Animal Health Pty Ltd) and 10mg/kg Ilium Xylazil-100 (Troy Laboratories), and pupils dilated with a drop of 1% Tropicamide (Alcon, Fort Worth, USA) applied on to the corneal surface. The AAV vectors were delivered using the UltraMicro Pump (UMP3) in combination with an RPE injection kit that consists of a SilFlex tubing and plastic needle holder and a NanoFil 10µl syringe (World Precision Instruments). For the subretinal injection, the mouse was positioned on its side under the dissecting microscope. Surgical scissors were used to gently cut through the connective tissue surrounding the injection site on the temporal, posterior part of the sclera. A small scleral incision was made using the tip of a disposable bevelled 26-gauge needle at the temporal side of the optic nerve. A blunt 35-gauge needle attached to the SilFlex tubing was inserted into the sclera at 45° from the eye. Once inside the eye, the angle of penetration was changed to 90° to avoid damage to the lens. When the tip of the needle reached the nasal subretinal space, the footswitch was used to release the required 1µl volume of vector at a rate of 250 nl/sec. To minimise backflow, the needle was kept in place for a further 5 seconds before it was removed from the eye. The final injected quantities per eye were 1e9 vg (AAV8 and Anc80 low dose), 3e9 vg (AAV8 and Anc80 medium dose) and 5e9 vg (AAV8 high dose). To confirm that the subretinal injection was successful, detachment in the inferior nasal (for the left eye) and the superior nasal (for the right eye) subretinal space was examined by OCT imaging using a Bioptigen OCT system (Leica) immediately after the subretinal injection. The animal then received a subcutaneous injection of Ilium Atipamezole (1mg/kg, Troy Laboratories) and topical antibiotic ointment (1% Chlorsig, Alcon Laboratories, Fort Worth, USA) was applied to the eye. The animals were returned to their cages and placed on a heated mat for recovery. Injected animals were also imaged using the Biotigen OCT at 1-week post-injection to confirm that the subretinal bleb had resolved. If a bleb or detachment was still present at this stage, the animal was excluded from further analysis.

### Electroretinogram (ERG) recordings

Retinal function was evaluated via full-field flash scotopic and photopic recordings performed on the Celeris full-field ERG system (Diagnosys LLC, Massachusetts, USA). Mice were dark-adapted overnight and handled under dim red light for scotopic readings. Mice were anesthetised with an intraperitoneal injection of 80mg/kg Ketamine (Ceva Animal Health Pty Ltd) and 10mg/kg Ilium Xylazil-100 (Troy Laboratories), and pupils dilated by application of a drop of 1% tropicamide to the cornea (Alcon, Fort Worth, USA). A drop of GenTeal (Alcon Laboratories, Fort Worth, USA) was also applied to the cornea and eye electrodes to ensure that moisture was retained and to act as a contact fluid. Animals were kept warm throughout the procedure with the in-built platform heater on the Celeris system. Scotopic and photopic recordings followed previously described protocols (85) and consisted of the following. For scotopic readings, single-flash intensities were obtained through 1ms flashes with intensities of 0.01, 0.1, 0.3, 1, 3, 10, and 25 cd.s.m-2. The time between consecutive flashes was 10 seconds, while the stimulus was repeated four times at 0.10Hz, with 60 seconds recovery time between different flash intensities. Following scotopic readings, mice were light-adapted for 10 minutes at 30 cd.s.m-2. A series of flashes on a background of 30 cd.m-2 were performed at 2 Hz and at intensities of 3 and 10 cd.s.m-2. The time between consecutive flashes was 0.5 seconds, and the stimulus was repeated 32 times. Flicker ERG responses were recorded at a pulse frequency of 10 and 30Hz, with a background of 30cd.m-2 and a pulse intensity of 3cd.s.m-2. Analysis of ERG waves, including extraction of the oscillatory potentials (OPs), was performed on the Espion V6 software (Diagnosys LLC, USA).

### Optomotor testing

Optomotor responses (OMR) were assessed using the automated OptoDrum system (Striatech, Germany) to measure the visual reflex to rotating stimuli. The OptoDrum is a closed box containing four screens that simulate a rotating cylinder of alternating white and black stripes. Mice were placed on a raised platform at the centre of the box, with a camera directly above. OMR movement was recorded and tracked using computer-controlled proprietary software and results automated. Mice were dark-adapted overnight and only handled under dim red light before scotopic recordings. Scotopic conditions were controlled during OMR testing by placing neutral density filter foils from the ScotopicKit (Striatech) on each screen inside the OptoDrum (ND4.8). This was followed by light adaptation for 2 hours before photopic recordings. Contrast sensitivity (%) and visual acuity (cycles/degree) were recorded under scotopic (2 mLux) and photopic (70 Lux) conditions, respectively.

### Immunohistochemistry and confocal imaging

Enucleated eyes from untreated and treated Kv8.2 KO, and wildtype (WT) mice were fixed for 30 minutes at room temperature in 4% paraformaldehyde (PFA, Electron Microscopy Science Inc., USA) in 1X Phosphate-buffered saline (PBS, Sigma-Aldrich). After fixation, the cornea, lens and sclera were removed, and eye cups placed back in 4% PFA for an additional 30 minutes at room temperature. Eye cups were then cryo-protected in a 20% sucrose solution overnight at 4°C, embedded in optimal cutting temperature (O.C.T compound; Tissue-Tek, Sakura Finetek, USA) media and cryosectioned on the sagittal plane at 12µm using a Leica (CM 3050S) cryostat. Retinal sections were rehydrated with 1X phosphate buffered saline (PBS) for 20 minutes at room temperature (RT) before blocking solution was applied for 1 hour at RT. Blocking solution contained 0.5% Triton X 100 (ThermoFisher, USA), 1% Bovine Serum Albumin (Bovogen Biologicals Pty Ltd., Australia), and 10% Normal Goat Serum (Sigma), diluted in 1X PBS. Primary antibodies (1:100 anti-Kv8.2 #75-014; 1:1000 anti-Kv2.1 #75-435, Neuromab Antibodies Inc) were applied overnight at 4°C, while secondary antibodies (1:500 Alexa Fluor^TM^, ThermoFisher. USA) were applied for 2 hours at RT. Following the application of primary and secondary antibodies, slides were incubated for 5 minutes with DAPI (4’, 6-diamidino-2-phenylindole, 0.5ug/mL in 1X PBS) and mounted using fluorescence mounting medium (Dako, USA). Negative controls were included in each batch by omitting the primary antibody. For Iba-1 staining for microglia quantification on wholemounts, eyes were enucleated and fixed in 4% PFA overnight at 4°C. After fixation retinas were then incubated in 10% normal goat serum, 3% Triton X-100, 1% BSA in 1X PBS for 1 hour at RT. Primary antibody incubation was carried out overnight at 4°C (1:500, FUJIFILM Wako, 019-19741). The retinas were then washed with 1X PBS and labelled with secondary antibody for 2 hours at RT at a 1:500 dilution in block solution. Confocal images were acquired on Nikon A1Si confocal microscope located at the UWA Harry Perkins Centre for Microscopy, Characterization and Analysis. For quantification of Iba-1 staining, confocal z stacks of vertical sections of retina labelled for Iba-1 and DAPI were acquired and quantified using an automated count in ImageJ (Fiji). In some cases, adjustments to image brightness and contrast were made with ImageJ (Fiji).

### RNA extraction and real-time quantitative polymerase chain reaction (qPCR)

Total RNA was extracted from whole retinas using TriReagent (Sigma-Aldrich) as per manufacturer’s instructions. Reverse transcription was performed using QuantiTect Reverse Transcription Kit (Qiagen) as per manufacturer’s instructions. Quantitative Real Time PCR (qPCR) was performed on a Bio-Rad CFX Connect Real-Time System using Taqman Fast Advanced mastermix (ThermoFisher) with the following TaqMan™ Gene Expression Assays (FAM): APYMN4P (codopt*KCNV2*), mouse *Kcnv2* (Mm00807577_m1) and mouse *Gapdh* (Mm99999915_g1). Gene expression was normalised to moue *Gapdh* and relative expression calculated using the ΔΔCt method.

### Cell dissociation for immune flow cytometry

Single-cell suspensions of retinal tissue were prepared as previously described(28). Briefly, eyes were dissected to separate the retina from the choroid/sclera. Retinas from both eyes were pooled for each mouse before the tissue was manually homogenised and digested in a mixture of 10 µg/ml Liberase (Roche, Basel, Switzerland) and 10 µg/ml DNAse I (Sigma, USA) in 1X PBS for 40 minutes at 37°C. The resulting single-cell preparations were stained with antibodies specific for CD45 (30F11), CD11b (M1/70), CD3 (145-2C11), CD4 (RM4-5), CD8 (53-6.7), NKp1.1 (PK136), CD11c (HL3), CD19 (6D5), CD64 (X54-5/7.1), F/480 (BM8), MHC-II (M5/114), Ly6C (AL.21), Ly6G (1AB). Antibodies were obtained from BD Biosciences (San Jose, CA, USA), BioLegend (San Diego, CA, USA), or eBioscience (San Diego, CA, USA). Fixable viability stain 620 (BD Biosciences) was used for live/dead discrimination. Samples were analysed using an LSRFortessa X-20 instrument (BD Biosciences). All data analysis was performed using the FlowJo software package (FlowJo, LLC, Ashland, OR, USA) and gating strategies followed the same steps as shown in(28).

### Single-cell RNA sequencing and bioinformatics

Preparation of single-cell suspensions of retinal cells was performed as described below using the solutions from the Worthington Biochemical Papain Dissociation System (Worthington Biochemical, USA). Enucleated eyes were dissected fresh in cold Hank’s balanced salt solution (HBSS) with Ca^2+^ and Mg^2+^ to remove the retina. Each individual retina was added to digestion solution pre-heated at 37°C with 20 units/ml papain and 0.005% DNase in EBSS. The sample was then incubated at 37°C for 15 minutes. After incubation, the digestion solution was removed and 600μl of inactivation solution (ovomucoid protease inhibitor [OMI] at 20 mg/ml; DNase 50 units/sample; 10% trehalose; 1% fetal calf serum; 0.4% bovine serum albumin) at RT was added to each sample, followed by gentle trituration using a glass pipette. 600μl of OMI was added, the samples centrifuged at 1000× rpm for 2 minutes and the supernatant was gently removed. 600μl of the resuspension solution (10% FCS; 0.4% BSA in EBSS) was added to the sample which was resuspended, passed through a 40µm cell strainer and kept on ice prior to processing for RNASeq. Individual retinal samples from four AAV8.KV-HD treated, two Kv8.2 KO untreated and two wildtype control animals were analysed.

Single cell libraries were created using the Single Cell 3’ Reagent Kit v3.1 (10X Genomics, California, USA) as per the manufacturer’s instructions. Briefly, individual cells were barcoded using gel beads-in-emulsion (GEMs) by adding the single cell suspensions, GEMs and partitioning oil loaded onto a Chromium NextGEM Chip G (10X Genomics, California USA). An estimated 20,000 cells were loaded to each channel to capture a recovery rate of approximately 10,000 cells. GEMs were then reverse transcribed for cDNA synthesis and samples were cleaned up using silane magnetic beads to purify the cDNA from the reaction mixture. Barcoded cDNA was amplified using PCR outlined in the Single Cell 3’ Reagent Kit v3.1, concentrations were detected by Qubit (Invitrogen, Massachusetts, USA), and sizing detected by Tapestation 4200 (Agilent, California, USA). Following these quality checks, cDNA was fragmented with sample indexes P5, P7, i7, i5, and TruSeq Read 2 added via end repair, A-tailing, adaptor ligation, and PCR. Gene expression libraries were sequenced on the NovaSeq 6000 Illumina platform (Illumina, California, USA). Expected sequencing depths were 18,000-20,000 reads per cell.

Quality of sequencing was checked on Illumina’s Sequence Analysis Viewer software and reads were demultiplexed using the bcl2fastq program using undetermined reads as an indicator of quality. FastQC and MultiQC were run and quality of reads was assessed based on adapter contamination, duplication rate, read diversity and sequencing error. Cell Ranger v7.01 was used to align reads to the mm10 reference genome and quantify gene expression of individual cells. The *KCNV2* codon-optimised human sequence was added to the reference genome to allow quantification of *KCNV2* expression. The raw and processed data has been uploaded to Gene Expression Omnibus (GEO) under accession number GSE311178.

The raw gene expression matrix was read into R version 4.3.2(86) and empty droplets were identified and removed using the DropletUtils package(87). Further quality control (QC) checks were performed using scater(88) to remove low quality cells that had less than 500 total reads or less than 200 detected genes. miQC(89) was used to fit a one dimensional mixture model and remove cells with a high proportion of reads mapping to the mitochondrial genome. Cells passing QC checks were subjected to further preprocessing and downstream analyses using Seurat v4.4.0(90). Firstly, genes without detectable expression in at least five cells were filtered out and then the data was normalised using SCTransform v2(91). Samples were integrated to correct for technical variation by selecting the top 3,000 most variable features and using these to identify anchors between cells across samples that are used to align all samples into a shared space. The integrated data was used to perform dimensional reduction (PCA & UMAP) and cell clustering. Clusters of cell types were annotated by examining expression of published marker genes from previous retinal tissue studies(56–59). Cells were deemed to have received gene therapy if they had detectable expression of *KCNV2*.The MAST statistical framework(92) was used within the Seurat FindMarkers() function to identify genes with 1) a log fold-change threshold of at least 0.18, 2) detection in at least 10% of cells in either group being tested and 3) at least three cells in either of the groups being tested. Biological replicates were accounted for as latent variables and genes with Bonferroni adjusted *p*-values less than 0.05 were called as differentially expressed. Enrichment and plotting of GO biological processes associated with the differentially expressed genes was performed using clusterProfiler(93).

### Proteomics

#### Sample processing and raw data preliminary analysis

Enucleated eyes were dissected in cold Phosphate-buffered saline, and the retina was isolated and placed into RIPA lysis buffer and supplemented with Halt™ Protease & Phosphatase Inhibitor Cocktail (ThermoFisher). The samples were sonicated for 10 seconds at an amplitude of 10 followed by centrifugation at 13000g for 15 minutes at 4°C. The supernatant was collected, and the total protein concentration was measured using Bio-Rad RC DC™ Protein Assay, as per manufacturer’s instructions. Retina solubilised proteins were reduced using 5 mM dithiothreitol and alkylated using 10 mM iodoacetamide. Proteins (150 µg) were initially digested at room temperature overnight using a 1:100 enzyme-to-protein ratio using Lys-C (Wako, Japan), followed by digestion with Trypsin (Promega, Madison, WI) for at least 4 hours at 37°C also at a 1:100 enzyme-to-protein ratio. Resultant peptides were acidified with 1% trifluoroacetic acid and purified using styrene divinylbenzene - reverse-phase sulfonate (Empore) stage tips. The proteome was identified on a Tandem Mass Tag (TMT) platform (Progenetech, Sydney, Australia).

#### Tandem Mass Tag (TMT) labelling

Three independent 10 plex TMT experiments were carried out. Ten independent PSEN1-pathogenic variant RPE and ten CRISPR edited isogenic controls were analysed for cell lysates; whilst five of each were analysed for upper chamber conditioned media to examine secreted protein. Briefly, dried peptides from each sample were resuspended in 100 mM HEPES (pH 8.2) buffer and peptide concentration measured using the MicroBCA protein assay kit. Sixty micrograms of peptide from each sample was subjected to TMT labelling with 0.8 mg of reagent per tube. Labelling was carried out at room temperature for 1 h with continuous vortexing. To quench any remaining TMT reagent and reverse the tyrosine labelling, 8 µl of 5% hydroxylamine was added to each tube, followed by vortexing and incubation for 15 min at room temperature. For each of the respective ten plex experiments, the ten labelled samples were combined, and then dried down by vacuum centrifugation. Prior to High-pH reversed-phase fractionation, the digested and TMT-labelled peptide samples were cleaned using a reverse-phase C18 clean-up column (Sep-pak, Waters) and dried in vacuum centrifuge. All labelled samples were combined to make a pooled sample. Peptides were desalted on a Strata-X 33 μm polymeric reversed phase column (Phenomenex). The combined samples were dissolved in loading buffer and separated by High pH on an Agilent 1100 HPLC system using a Zorbax C18 column (2.1 x 150 mm). Peptides were eluted with a linear gradient of 20mM ammonium formate, 2% ACN to 20mM ammonium formate, 90% ACN at 0.2ml/min. The 95 fractions were concatenated into 12 fractions and dried down.

#### LC-ESI-MS/MS data acquisition

Fractions were analysed by electrospray ionisation mass spectrometry using a Thermo UltiMate 3000 nanoflow UHPLC system (Thermo Scientific) coupled to a Orbitrap Exploris 480 mass spectrometer (Thermo Scientific). Peptides were loaded onto an Reprosil-pur C18 LC Column, 1.9μm particle size x 150mm and separated with a linear gradient of water/acetonitrile/0.1% formic acid (v/v). The initial analyses were carried out on the NCRIS (National Collaborative Research Infrastructure Strategy) enabled Bioplatforms Australia infrastructure at the Western Australian Proteomics Facility, located at the Harry Perkins Institute for Medical Research.

#### Proteomic data analysis

Spectra were converted to mzXML via MSconvert v3.0. Database searching included all entries from the Mus musculus. UniProt reference Database (downloaded: June 2024). The database was concatenated with one composed of all protein sequences for that database in the reversed order. Searches were performed using a 50-ppm precursor ion tolerance for total protein-level profiling. The product ion tolerance was set to 0.2 Da. These wide mass tolerance windows were chosen to maximise sensitivity in conjunction with Comet searches and linear discriminant analysis. TMT tags on lysine residues and peptide N-termini (+ 229.163 Da for TMT) and carbamidomethylation of cysteine residues (+ 57.021 Da) were set as static modifications, while oxidation of methionine residues (+ 15.995 Da) was set as a variable modification. Peptide-spectrum matches (PSMs) were adjusted to a 1% false discovery rate (FDR). PSM filtering was performed using a linear discriminant analysis, as described previously and then assembled further to a final protein-level FDR of 1%, using the Picked FDR method. An isolation purity of at least 0.7 (70%) in the MS1 isolation window was used for samples. For each protein, the filtered peptide TMT SN values were summed to create protein quantifications. To control for different total protein loading within a TMT experiment, the summed protein quantities of each channel were adjusted to be equal within the experiment. Proteins were quantified by summing reporter ion counts across all matching PSMs, also as described previously. Reporter ion intensities were adjusted to correct for the isotopic impurities of the different TMT reagents according to manufacturer specifications. Finally, each protein abundance measurement was scaled, such that the summed signal-to-noise for that protein across all channels equaled 100, thereby generating a relative abundance (RA) measurement.

### Retinal organoids

#### Generation of human iPSCs from PBMCs

Following informed consent, peripheral blood was taken from an individual with confirmed Cone dystrophy with Supernormal Rod Electroretinogram inherited retinal degeneration. This study was approved by the Sydney Children’s Hospitals Network Human Research Ethics Committee, HREC/17/SCHN/323. The PBMC isolation and their reprogramming were performed as previously described(94).

#### Human iPSC maintenance and retinal differentiation culture

The hiPSC lines were maintained on feeder free conditions in Essential 8 medium and Geltrex™ (ThermoFisher) coated 6-well plates. For retinal neuroepithelial differentiation hiPSCs were maintained until cells reached 90-95 % confluency, then medium without FGF (Essential 6 medium) was added to the cultures for two days, followed by a neural induction for up to 6 weeks in proneural induction medium (Advanced DMEM/F12, 1X MEM non-essential amino acids, 1X N2 Supplement, 100 mM glutamine; ThermoFisher). Lightly pigmented islands of retinal pigmented epithelium (RPE) appeared as early as week 3 in culture. Optic vesicles were formed within the RPE regions between weeks 4 and 6, these neural retinal vesicles were manually excised with 18G needles and kept individually in low binding 96 well plates in retinal differentiation medium (DMEM/F12 with high glucose, 2% B27 minus vitamin A). At 6 weeks of differentiation, the retinal differentiation medium was supplemented with 10 % FBS, 100 uM taurine and 2 mM glutamax. From 10 weeks, vesicles were transferred to low binding 24 well plates (5 vesicles/well) and was cultured in maturation medium (Advanced DMEM, 10% FBS, 2% B27 minus vitamin A, 1X Glutamax, 100 uM Taurine supplemented with 1 uM retinoic acid (RA). At 12 weeks of differentiation the FBS was removed from medium and supplemented with 1X N2 supplement and 0.5 µM RA concentration. Maintenance cultures of hiPSCs were fed daily and differentiation cultures were fed every 2 days. At 26 weeks, hiPSC-derived retinal organoids were collected for downstream experiments.

#### Viral transduction of Retinal Organoids

At 20 weeks of differentiation, individual retinal organoids were transferred to low binding 24 well plates (Costar, Corning). AAV2/7m8.RK.hKCNV2 viral vector (1x10E11 vg/organoid) was added to a total volume of 375 μL using fresh medium used to culture the retinal organoids. The retinal organoids were incubated at 37 °C overnight before adding another topping up with fresh medium. Cultures were fed as normal until week 26 of differentiation.

#### Human iPSC-Retinal Organoid Immunohistochemistry

HiPSC-RO were collected and fixed for 45 minutes in 4% paraformaldehyde and refrigerated overnight in 20% sucrose prior to embedding in OCT. Cryosections (14 µM) were blocked in 5% goat serum and 1% bovine serum albumin in PBS for 2 hours. Primary antibody was incubated overnight at 4°C. Sections were incubated with secondary antibody for 2 hrs at room temperature, washed and counter-stained with DAPI (Sigma). Alexa fluor 488 secondary antibody (Invitrogen-Molecular Probes) was used at a 1:500 dilution. Primary antibodies Anti-KCVN2 (Sigma; HPA031131; 1:100), Monoclonal Anti-Opsin (Merck; O4886; 1:1000) and Peripherin-2 (Merck; MABN293; 1:800). Sections were imaged on the Leica Stellaris 8 confocal microscope and images were processed using LASX software.

#### Western Blots

HiPSC-retinal organoids were homogenised and lysed in RIPA buffer and protease inhibitor cocktail. The samples were centrifuged at 17000g for 15 minutes. The protein concentration was measured using Pierce™BCA Protein Assay Kits; for each sample 20 µg of protein was incubated (37°C for 30 minutes) in NuPAGE™ LDS sample buffer and sample reducing agent. The samples were run on NuPAGE™ 4 to 12%, Bis-Tris, 1.0–1.5mm, Mini Protein Gel at 150 volts for 50 minutes. The gel was transferred to a 0.45 μm nitrocellulose membrane at 30 volts for 90 minutes in 20% v/v methanol and NuPAGE™ Transfer buffer. The membrane was blocked with 5% BSA, 1XTBS (10X TBS: 24g Tris and 88g NaCl in 1L water pH 7.6) for 2 hours, then incubated overnight at 4°C with anti-KCVN2 (Sigma Aldrich: HPA031131) and anti-beta actin (Abcam: ab8225). The blots were washed in 1XTBS and incubated with IRDye® 680RD Goat anti-Rabbit IgG Secondary Antibody and IRDye® 800CW Goat anti-Mouse IgG Secondary Antibody in the dark for 1 hour before imaging using Chemidoc Touch Gel Imaging System (Bio-Rad).

#### Statistical analysis

All means are presented ± SD (standard deviation), unless otherwise stated; N, number of independent experiments performed; For quantification assessment, statistical analysis is based on at least three independent experiments. Statistical significance was assessed using Mann-Whitney t-test or ANOVA in Graphpad Prism 6 software and denoted as *p*.

### Statistical Analysis

Statistical analysis was performed using the PRISM software (GraphPad; San Diego, USA) and Excel (Microsoft; Washington, USA). Either t-test or ANOVA was used to determine statistical significance (*P*<0.05). Further statistical analysis for scRNASeq and proteomics analysis are provided under their specific sections in the Methods.

### Animals and study approvals

All animal use was approved by the Institutional Animal Ethics Committee of the Harry Perkins Medical Research Institute (AE216) and was in accordance with the Australian Code for the Care and Use of Animals for Scientific Purposes and the ARVO Statement for the Use of Animals in Ophthalmic and Vision Research. Mice from both sexes were used and all mice were group-housed in a climate-controlled facility on a 12-hour light/dark cycle with *ad libitum* access to food and water. The Kv8.2 KO line was generated at the Wellcome Trust Sanger. A full description of how this line was generated can be found in (33, 95). Homozygote carriers were inter-crossed to generate homozygote animals for the experiments described in this study. Wildtype C57BL/6J mice were purchased from the Animal Resource Centre (ARC, now under Ozgene, Australia).

## Authors contributions

LSC, DMH, RR designed the research study. RR conducted experiments, acquired data, analysed data, and prepared the first draft of figures. XRL, PIFC, AAB, ALM, YB, REJ, DA, VV, JAP, MMizraei, MMMangala, EOW, RVJ, AGC acquired and/or analysed data. LSC and DMH wrote the manuscript and prepared final figures. All authors reviewed the manuscript.

## Acknowledgments

The authors acknowledge the facilities, and the scientific and technical assistance of Microscopy Australia at the Centre for Microscopy, Characterisation & Analysis, The University of Western Australia, a facility funded by the University, State, and Commonwealth Governments.

## References

1. Georgiou M, Fujinami K, Vincent A, Nasser F, Khateb S, Vargas ME, et al. KCNV2-Associated Retinopathy: Detailed Retinal Phenotype and Structural Endpoints-KCNV2 Study Group Report 2. Am J Ophthalmol. 2021;230:1–11.

2. Georgiou M, Robson AG, Fujinami K, Leo SM, Vincent A, Nasser F, et al. KCNV2-Associated Retinopathy: Genetics, Electrophysiology, and Clinical Course-KCNV2 Study Group Report 1. Am J Ophthalmol. 2021;225:95–107.

3. Robson AG, Webster AR, Michaelides M, Downes SM, Cowing JA, Hunt DM, et al. “Cone dystrophy with supernormal rod electroretinogram”: a comprehensive genotype/phenotype study including fundus autofluorescence and extensive electrophysiology. Retina. 2010;30(1):51–62.

4. Thiagalingam S, McGee TL, Weleber RG, Sandberg MA, Trzupek KM, Berson EL, and Dryja TP. Novel mutations in the KCNV2 gene in patients with cone dystrophy and a supernormal rod electroretinogram. Ophthalmic Genet. 2007;28(3):135–42.

5. Wissinger B, Dangel S, Jagle H, Hansen L, Baumann B, Rudolph G, et al. Cone dystrophy with supernormal rod response is strictly associated with mutations in KCNV2. Investigative ophthalmology & visual science. 2008;49(2):751–7.

6. Wu TH, Ting TD, Okajima TI, Pepperberg DR, Ho YK, Ripps H, and Naash MI. Opsin localization and rhodopsin photochemistry in a transgenic mouse model of retinitis pigmentosa. Neuroscience. 1998;87(3):709–17.

7. Abdelkader E, Yasir ZH, Khan AM, Raddadi O, Khandekar R, Alateeq N, et al. Analysis of retinal structure and function in cone dystrophy with supernormal rod response. Doc Ophthalmol. 2020;141(1):23–32.

8. Greenwald SH, Kuchenbecker JA, Rowlan JS, Neitz J, and Neitz M. Role of a Dual Splicing and Amino Acid Code in Myopia, Cone Dysfunction and Cone Dystrophy Associated with L/M Opsin Interchange Mutations. Transl Vis Sci Technol. 2017;6(3):2.

9. Guimaraes TAC, Georgiou M, Robson AG, and Michaelides M. KCNV2 retinopathy: clinical features, molecular genetics and directions for future therapy. Ophthalmic Genet. 2020;41(3):208–15.

10. Fujinami K, Tsunoda K, Nakamura N, Kato Y, Noda T, Shinoda K, et al. Molecular characteristics of four Japanese cases with KCNV2 retinopathy: report of novel disease-causing variants. Mol Vis. 2013;19:1580–90.

11. Vincent A, Wright T, Garcia-Sanchez Y, Kisilak M, Campbell M, Westall C, and Heon E. Phenotypic characteristics including in vivo cone photoreceptor mosaic in KCNV2-related “cone dystrophy with supernormal rod electroretinogram”. Investigative ophthalmology & visual science. 2013;54(1):898–908.

12. Gouras P, Eggers HM, and MacKay CJ. Cone dystrophy, nyctalopia, and supernormal rod responses. A new retinal degeneration. Arch Ophthalmol. 1983;101(5):718–24.

13. Hood DC, Cideciyan AV, Halevy DA, and Jacobson SG. Sites of disease action in a retinal dystrophy with supernormal and delayed rod electroretinogram b-waves. Vision Res. 1996;36(6):889–901.

14. Kato M, Kobayashi R, and Watanabe I. Cone dysfunction and supernormal scotopic electroretinogram with a high-intensity stimulus. A report of three cases. Doc Ophthalmol. 1993;84(1):71–81.

15. Michaelides M, Holder GE, Webster AR, Hunt DM, Bird AC, Fitzke FW, et al. A detailed phenotypic study of “cone dystrophy with supernormal rod ERG”. Br J Ophthalmol. 2005;89(3):332–9.

16. Rosenberg T, and Simonsen SE. Retinal cone dysfunction of supernormal rod ERG type. Five new cases. Acta Ophthalmol (Copenh). 1993;71(2):246–55.

17. Wu H, Cowing JA, Michaelides M, Wilkie SE, Jeffery G, Jenkins SA, et al. Mutations in the gene *KCNV2* encoding a voltage-gated potassium channel subunit cause “cone dystrophy with a supernormal rod electroretinogram” in humans. Amer J Hum Genet. 2006;79:574–9.

18. In: National Center for Biotechnology I ed. Bethesda, MD, USA: National Library of Medicine.

19. Wilson NH, and Key B. Neogenin interacts with RGMa and netrin-1 to guide axons within the embryonic vertebrate forebrain. Developmental biology. 2006;296(2):485–98.

20. Kang SW, Leclerc B, Kosonsiriluk S, Mauro LJ, Iwasawa A, and El Halawani ME. Melanopsin expression in dopamine-melatonin neurons of the premammillary nucleus of the hypothalamus and seasonal reproduction in birds. Neuroscience. 2010;170(1):200–13.

21. Ben Salah S, Kamei S, Senechal A, Lopez S, Bazalgette C, Eliaou CM, et al. Novel KCNV2 mutations in cone dystrophy with supernormal rod electroretinogram. Am J Ophthalmol. 2008;145(6):1099–106.

22. Sandberg MA, Miller S, and Berson EL. Rod electroretinograms in an elevated cyclic guanosine monophosphate-type human retinal degeneration. Comparison with retinitis pigmentosa. Investigative ophthalmology & visual science. 1990;31(11):2283–7.

23. Weng J, Mata NL, Azarian SM, Tzekov RT, Birch DG, and Travis GH. Insights into the function of Rim protein in photoreceptors and etiology of Stargardt’s disease from the phenotype in abcr knockout mice. Cell. 1999;98(1):13–23.

24. Smith KE, Wilkie SE, Tebbs-Warner JT, Jarvis BJ, Gallasch L, Stocker M, and Hunt DM. Functional analysis of missense mutations in Kv8.2 causing cone dystrophy with supernormal rod electroretinogram. J Biol Chem. 2012;287(52):43972–83.

25. Czirjak G, Toth ZE, and Enyedi P. Characterization of the heteromeric potassium channel formed by kv2.1 and the retinal subunit kv8.2 in Xenopus oocytes. J Neurophysiol. 2007;98(3):1213–22.

26. Gayet-Primo J, Yaeger DB, Khanjian RA, and Puthussery T. Heteromeric KV2/KV8.2 channels mediate delayed rectifier potassium currents in primate photoreceptors. The Journal of neuroscience : the official journal of the Society for Neuroscience. 2018;In press.

27. Hart NS, Mountford JK, Carvalho LS, Barth M, Voight V, Nerbonne JM, and Hunt DM. The role of the voltage-gated potassium channel proteins Kv8.2 and Kv2.1 in vision and retinal disease; insights from the study of mouse gene knock-out mutations. 2017.

28. Jiang X, Rashwan R, Voigt V, Nerbonne J, Hunt DM, and Carvalho LS. Molecular, Cellular and Functional Changes in the Retinas of Young Adult Mice Lacking the Voltage-Gated K+ Channel Subunits Kv8.2 and K2.1. International Journal of Molecular Sciences. 2021;22(9):4877.

29. Sakti DH, Cornish EE, Ali H, Retsas S, Raza M, Saakova N, et al. Natural history and biomarkers of KCNV2-associated retinopathy. Clinical & experimental ophthalmology. 2024;52(5):528–44.

30. Collison FT, Park JC, Fishman GA, Stone EM, and McAnany JJ. Two-color pupillometry in KCNV2 retinopathy. Doc Ophthalmol. 2019;139(1):11–20.

31. Trapani I, and Auricchio A. Has retinal gene therapy come of age? From bench to bedside and back to bench. Hum Mol Genet. 2019;28(R1):R108–R18.

32. Kaiser VM, and Gonzalez-Cordero A. Organoids - the future of pre-clinical development of AAV gene therapy for CNS disorders. Gene Ther. 2025.

33. Hart NS, Mountford JK, Voigt V, Fuller-Carter P, Barth M, Nerbonne JM, et al. The role of the voltage-gated potassium channel proteins Kv8.2 and Kv2.1 in vision and retinal disease: insights from the study of mouse gene knock-out mutations. eneuro. 2019:ENEURO.0032-19.2019.

34. Gayet-Primo J, Yaeger DB, Khanjian RA, and Puthussery T. Heteromeric KV2/KV8.2 Channels Mediate Delayed Rectifier Potassium Currents in Primate Photoreceptors. J Neurosci. 2018;38(14):3414–27.

35. Hart NS, Mountford JK, Voight V, Fuller-Carter P, Barth M, Nerbonne JM, et al. The role of the voltage-gated potassium channel proteins Kv8.2 and Kv2.1 in vision and retinal disease; insights from the study of mouse gene knock-out mutations. eNeuro. 2019;6(1).

36. Beech DJ, and Barnes S. Characterization of a voltage-gated K+ channel that accelerates the rod response to dim light Neuron. 1989;3(5):573–81.

37. Kurenny DE, and Barnes S. Proton modulation of M-like potassium current (IKx) in rod photoreceptors. Neurosci Lett. 1994;170(2):225–8.

38. Barrow AJ, and Wu SM. Complementary conductance changes by IKx and Ih contribute to membrane impedance stability during the rod light response. Channels (Austin). 2009;3(5):301–7.

39. Young JE, Vogt T, Gross KW, and Khani SC. A short, highly active photoreceptor-specific enhancer/promoter region upstream of the human rhodopsin kinase gene. Investigative ophthalmology & visual science. 2003;44(9):4076–85.

40. Fuller-Carter PI, Basiri H, Harvey AR, and Carvalho LS. Focused Update on AAV-Based Gene Therapy Clinical Trials for Inherited Retinal Degeneration. BioDrugs. 2020;34(6):763–81.

41. Carvalho LS, Xiao R, Wassmer S, Langsdorf A, Zinn E, Pacouret S, et al. Synthetic adeno-associated viral vector efficiently targets mouse and non-human primate retina in vivo. Hum Gene Ther. 2018.

42. Vandenberghe LH, Bell P, Maguire AM, Cearley CN, Xiao R, Calcedo R, et al. Dosage thresholds for AAV2 and AAV8 photoreceptor gene therapy in monkey. Sci Transl Med. 2011;3(88):88ra54.

43. Marangoni D, Bush RA, Zeng Y, Wei LL, Ziccardi L, Vijayasarathy C, et al. Ocular and systemic safety of a recombinant AAV8 vector for X-linked retinoschisis gene therapy: GLP studies in rabbits and Rs1-KO mice. Mol Ther Methods Clin Dev. 2016;5:16011.

44. Cukras C, Wiley HE, Jeffrey BG, Sen HN, Turriff A, Zeng Y, et al. Retinal AAV8-RS1 Gene Therapy for X-Linked Retinoschisis: Initial Findings from a Phase I/IIa Trial by Intravitreal Delivery. Mol Ther. 2018;26(9):2282–94.

45. Cehajic-Kapetanovic J, Xue K, Martinez-Fernandez de la Camara C, Nanda A, Davies A, Wood LJ, et al. Initial results from a first-in-human gene therapy trial on X-linked retinitis pigmentosa caused by mutations in RPGR. Nat Med. 2020;26(3):354–9.

46. Fischer MD, Michalakis S, Wilhelm B, Zobor D, Muehlfriedel R, Kohl S, et al. Safety and Vision Outcomes of Subretinal Gene Therapy Targeting Cone Photoreceptors in Achromatopsia: A Nonrandomized Controlled Trial. JAMA Ophthalmol. 2020.

47. Perlman I. In: Kolb H, Fernandez E, and Nelson R eds. Webvision: The Organization of the Retina and Visual System. Salt Lake City (UT); 1995.

48. Wu H, Cowing JA, Michaelides M, Wilkie SE, Jeffery G, Jenkins SA, et al. Mutations in the gene KCNV2 encoding a voltage-gated potassium channel subunit cause “cone dystrophy with supernormal rod electroretinogram” in humans. Am J Hum Genet. 2006;79(3):574–9.

49. Sergouniotis PI, Holder GE, Robson AG, Michaelides M, Webster AR, and Moore AT. High-resolution optical coherence tomography imaging in KCNV2 retinopathy. Br J Ophthalmol. 2012;96(2):213–7.

50. Wachtmeister L, and Dowling JE. The oscillatory potentials of the mudpuppy retina. Investigative ophthalmology & visual science. 1978;17(12):1176–88.

51. Wachtmeister L, and Hahn I. Spatial properties of the oscillatory potentials of the frog electroretinogram in relation to state of adaptation. Acta Ophthalmol (Copenh). 1987;65(6):724–30.

52. Algvere P, Wachtmeister L, and Westbeck S. ON THE OSCILLATORY POTENTIALS OF THE HUMAN ELECTRORETINOGRAM IN LIGHT AND DARK ADAPTATION: I. Thresholds and relation to stimulus intensity on adaptation to short flashes of light. A Fourier analysis. Acta ophthalmologica. 1972;50(5):737–59.

53. Wachtmeister L. On the oscillatory potentials of the human electroretinogram in light and dark adaptation: III. Thresholds and relation to stimulus intensity on adaptation to background light. Acta Ophthalmologica. 1973;51(1):95–113.

54. Speros P, and Price J. Oscillatory potentials. History, techniques and potential use in the evaluation of disturbances of retinal circulation. Survey of Ophthalmology. 1981;25(4):237–52.

55. King-Smith PE, Loffing D, and Jones R. Rod and cone ERGs and their oscillatory potentials. Investigative ophthalmology & visual science. 1986;27(2):270–3.

56. Macosko EZ, Basu A, Satija R, Nemesh J, Shekhar K, Goldman M, et al. Highly Parallel Genome-wide Expression Profiling of Individual Cells Using Nanoliter Droplets. Cell. 2015;161(5):1202–14.

57. Karademir D, Todorova V, Ebner LJA, Samardzija M, and Grimm C. Single-cell RNA sequencing of the retina in a model of retinitis pigmentosa reveals early responses to degeneration in rods and cones. BMC Biol. 2022;20(1):86.

58. Clark BS, Stein-O’Brien GL, Shiau F, Cannon GH, Davis-Marcisak E, Sherman T, et al. Single-Cell RNA-Seq Analysis of Retinal Development Identifies NFI Factors as Regulating Mitotic Exit and Late-Born Cell Specification. Neuron. 2019;102(6):1111–26 e5.

59. Shekhar K, Lapan SW, Whitney IE, Tran NM, Macosko EZ, Kowalczyk M, et al. Comprehensive Classification of Retinal Bipolar Neurons by Single-Cell Transcriptomics. Cell. 2016;166(5):1308–23 e30.

60. Fadl BR, Brodie SA, Malasky M, Boland JF, Kelly MC, Kelley MW, et al. An optimized protocol for retina single-cell RNA sequencing. Mol Vis. 2020;26:705–17.

61. Lang F, and Shumilina E. Regulation of ion channels by the serum- and glucocorticoid-inducible kinase SGK1. FASEB J. 2013;27(1):3–12.

62. Weh E, Lutrzykowska Z, Smith A, Hager H, Pawar M, Wubben TJ, and Besirli CG. Hexokinase 2 is dispensable for photoreceptor development but is required for survival during aging and outer retinal stress. Cell Death Dis. 2020;11(6):422.

63. Agrawal SA, Burgoyne T, Eblimit A, Bellingham J, Parfitt DA, Lane A, et al. REEP6 deficiency leads to retinal degeneration through disruption of ER homeostasis and protein trafficking. Hum Mol Genet. 2017;26(14):2667–77.

64. Tomita Y, Qiu C, Bull E, Allen W, Kotoda Y, Talukdar S, et al. Muller glial responses compensate for degenerating photoreceptors in retinitis pigmentosa. Exp Mol Med. 2021;53(11):1748–58.

65. Hawes BE, Touhara K, Kurose H, Lefkowitz RJ, and Inglese J. Determination of the G beta gamma-binding domain of phosducin. A regulatable modulator of G beta gamma signaling. J Biol Chem. 1994;269(47):29825–30.

66. Gonzalez-Cordero A, Goh D, Kruczek K, Naeem A, Fernando M, Kleine Holthaus SM, et al. Assessment of AAV Vector Tropisms for Mouse and Human Pluripotent Stem Cell-Derived RPE and Photoreceptor Cells. Hum Gene Ther. 2018.

67. Sun X, Pawlyk B, Xu X, Liu X, Bulgakov OV, Adamian M, et al. Gene therapy with a promoter targeting both rods and cones rescues retinal degeneration caused by AIPL1 mutations. Gene therapy. 2010;17(1):117–31.

68. Boye SE, Boye SL, Pang J, Ryals R, Everhart D, Umino Y, et al. Functional and behavioral restoration of vision by gene therapy in the guanylate cyclase-1 (GC1) knockout mouse. PLoS One. 2010;5(6):e11306.

69. Mihelec M, Pearson RA, Robbie SJ, Buch PK, Azam SA, Bainbridge JW, et al. Long-term preservation of cones and improvement in visual function following gene therapy in a mouse model of leber congenital amaurosis caused by guanylate cyclase-1 deficiency. Human gene therapy. 2011;22(10):1179–90.

70. Boye SE, Alexander JJ, Boye SL, Witherspoon CD, Sandefer KJ, Conlon TJ, et al. The human rhodopsin kinase promoter in an AAV5 vector confers rod- and cone-specific expression in the primate retina. Hum Gene Ther. 2012;23(10):1101–15.

71. Boye SL, Peshenko IV, Huang WC, Min SH, McDoom I, Kay CN, et al. AAV-mediated gene therapy in the guanylate cyclase (RetGC1/RetGC2) double knockout mouse model of Leber congenital amaurosis. Hum Gene Ther. 2013;24(2):189–202.

72. Hsu Y, Bhattarai S, Thompson JM, Mahoney A, Thomas J, Mayer SK, et al. Subretinal gene therapy delays vision loss in a Bardet-Biedl Syndrome type 10 mouse model. Mol Ther Nucleic Acids. 2023;31:164–81.

73. Storchi R, Rodgers J, Gracey M, Martial FP, Wynne J, Ryan S, et al. Measuring vision using innate behaviours in mice with intact and impaired retina function. Sci Rep. 2019;9(1):10396.

74. Farahbakhsh M, Anderson EJ, Maimon-Mor RO, Rider A, Greenwood JA, Hirji N, et al. A demonstration of cone function plasticity after gene therapy in achromatopsia. Brain. 2022;145(11):3803–15.

75. Michaelides M, Hirji N, Wong SC, Besirli CG, Zaman S, Kumaran N, et al. First-in-Human Gene Therapy Trial of AAV8-hCARp.hCNGB3 in Adults and Children With CNGB3-associated Achromatopsia. Am J Ophthalmol. 2023;253:243–51.

76. Carvalho LS, Xu J, Pearson RA, Smith AJ, Bainbridge JW, Morris LM, et al. Long-term and age-dependent restoration of visual function in a mouse model of CNGB3-associated achromatopsia following gene therapy. Hum Mol Genet. 2011;20(16):3161–75.

77. Irigoyen C, Amenabar Alonso A, Sanchez-Molina J, Rodriguez-Hidalgo M, Lara-Lopez A, and Ruiz-Ederra J. Subretinal Injection Techniques for Retinal Disease: A Review. J Clin Med. 2022;11(16).

78. Britten-Jones AC, Jin R, Gocuk SA, Cichello E, O’Hare F, Hickey DG, et al. The safety and efficacy of gene therapy treatment for monogenic retinal and optic nerve diseases: A systematic review. Genet Med. 2022;24(3):521–34.

79. Khanani AM, Aziz AA, Khanani ZA, Khan H, Mojumder O, Sulahria H, et al. Subretinal Gene Therapy for Treatment of Retinal and Choroidal Vascular Diseases. Am J Ophthalmol. 2024.

80. Bainbridge JW, Smith AJ, Barker SS, Robbie S, Henderson R, Balaggan K, et al. Effect of gene therapy on visual function in Leber’s congenital amaurosis. The New England journal of medicine. 2008;358(21):2231–9.

81. Hauswirth WW, Aleman TS, Kaushal S, Cideciyan AV, Schwartz SB, Wang L, et al. Treatment of leber congenital amaurosis due to RPE65 mutations by ocular subretinal injection of adeno-associated virus gene vector: short-term results of a phase I trial. Human gene therapy. 2008;19(10):979–90.

82. Maguire AM, Simonelli F, Pierce EA, Pugh EN, Jr., Mingozzi F, Bennicelli J, et al. Safety and efficacy of gene transfer for Leber’s congenital amaurosis. The New England journal of medicine. 2008;358(21):2240–8.

83. Marshall LJ, Bailey J, Cassotta M, Herrmann K, and Pistollato F. Poor Translatability of Biomedical Research Using Animals - A Narrative Review. Altern Lab Anim. 2023;51(2):102–35.

84. Loeb JE, Cordier WS, Harris ME, Weitzman MD, and Hope TJ. Enhanced expression of transgenes from adeno-associated virus vectors with the woodchuck hepatitis virus posttranscriptional regulatory element: implications for gene therapy. Hum Gene Ther. 1999;10(14):2295–305.

85. Brunet AA, Fuller-Carter PI, Miller AL, Voigt V, Vasiliou S, Rashwan R, et al. Validating Fluorescent Chrnb4.EGFP Mouse Models for the Study of Cone Photoreceptor Degeneration. Transl Vis Sci Technol. 2020;9(9):28.

86. R Core Team. Foundation for Statistical Computing, Vienna, Austria; 2025.

87. Griffiths JA, Richard AC, Bach K, Lun ATL, and Marioni JC. Detection and removal of barcode swapping in single-cell RNA-seq data. Nat Commun. 2018;9(1):2667.

88. McCarthy DJ, Campbell KR, Lun AT, and Wills QF. Scater: pre-processing, quality control, normalization and visualization of single-cell RNA-seq data in R. Bioinformatics. 2017;33(8):1179–86.

89. Hippen AA, Falco MM, Weber LM, Erkan EP, Zhang K, Doherty JA, et al. miQC: An adaptive probabilistic framework for quality control of single-cell RNA-sequencing data. PLoS Comput Biol. 2021;17(8):e1009290.

90. Hao Y, Hao S, Andersen-Nissen E, Mauck WM, 3rd, Zheng S, Butler A, et al. Integrated analysis of multimodal single-cell data. Cell. 2021;184(13):3573–87 e29.

91. Choudhary S, and Satija R. Comparison and evaluation of statistical error models for scRNA-seq. Genome Biol. 2022;23(1):27.

92. Finak G, McDavid A, Yajima M, Deng J, Gersuk V, Shalek AK, et al. MAST: a flexible statistical framework for assessing transcriptional changes and characterizing heterogeneity in single-cell RNA sequencing data. Genome Biol. 2015;16:278.

93. Yu G, Wang LG, Han Y, and He QY. clusterProfiler: an R package for comparing biological themes among gene clusters. OMICS. 2012;16(5):284–7.

94. Ribeiro J, Procyk CA, West EL, O’Hara-Wright M, Martins MF, Khorasani MM, et al. Restoration of visual function in advanced disease after transplantation of purified human pluripotent stem cell-derived cone photoreceptors. Cell Rep. 2021;35(3):109022.

95. Perez-Riverol Y, Bandla C, Kundu DJ, Kamatchinathan S, Bai J, Hewapathirana S, et al. The PRIDE database at 20 years: 2025 update. Nucleic Acids Res. 2025;53(D1):D543–D53.

