## Supplementary data for "Pre-clinical validation of a novel AAV-mediated gene therapy for *KCNV2* retinopathy improves visual function in a mouse model and expression in patient organoids"

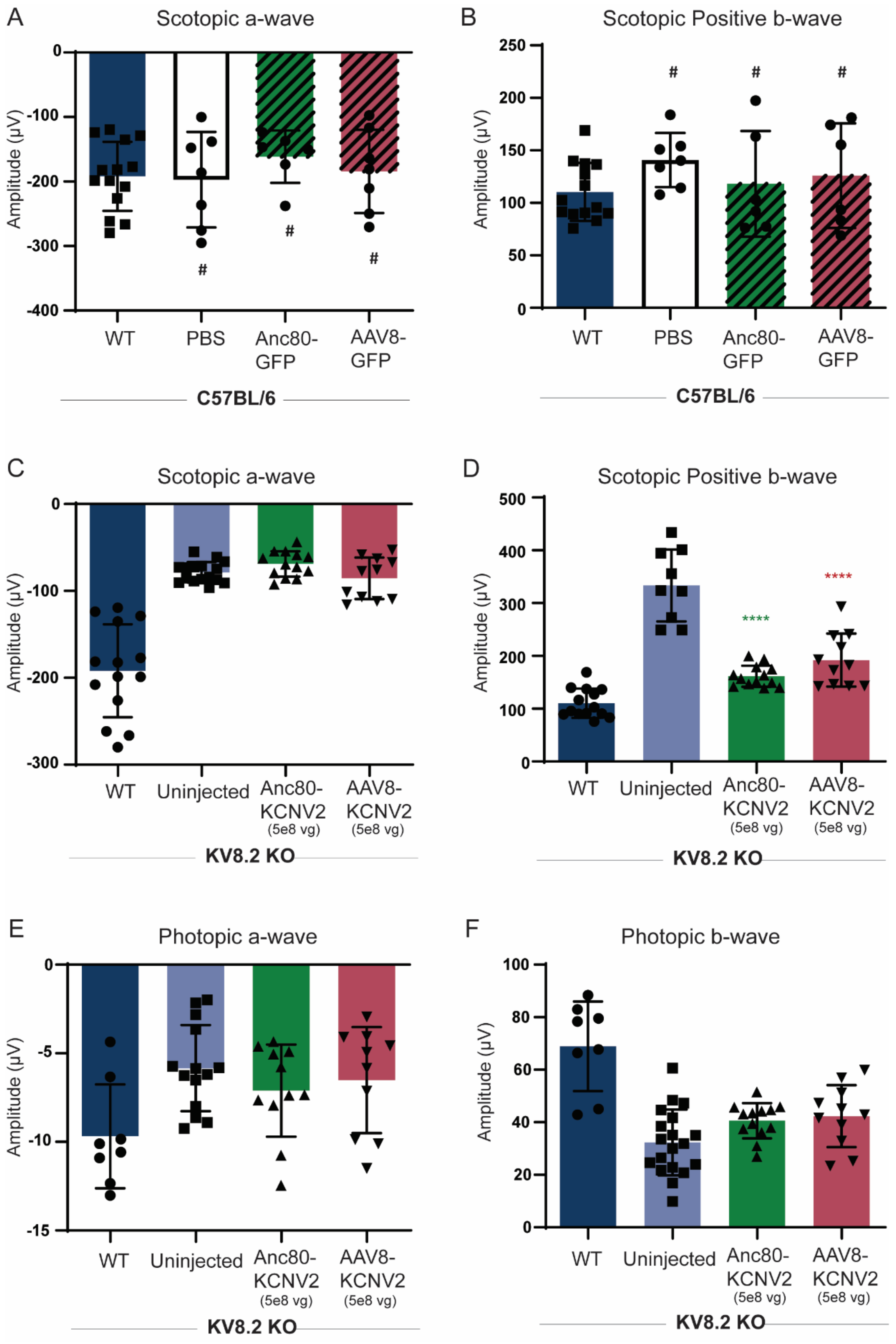


**Supplementary Figure 1. Dark-adapted and light-adapted electroretinogram (ERG) responses from AAV.GFP and low dose treated groups. (A-B)** Dark-adapted (scotopic) a-wave and positive b-wave ERG responses from wildtype C57BL/6 mice treated with either Anc80.GFP or AAV8.GFP vectors. **(C-D)** Dark-adapted (scotopic) a-wave and positive b-wave ERG responses from Kv8.2 KO mice treated with either Anc80.KCNV2 or AAV8.KCNV2 vectors at 5e8 vg. (E-F) Light-adapted (photopic) a-wave and b-wave ERG responses from Kv8.2 KO mice treated with either Anc80.KCNV2 or AAV8.KCNV2 vectors at 5e8 vg.

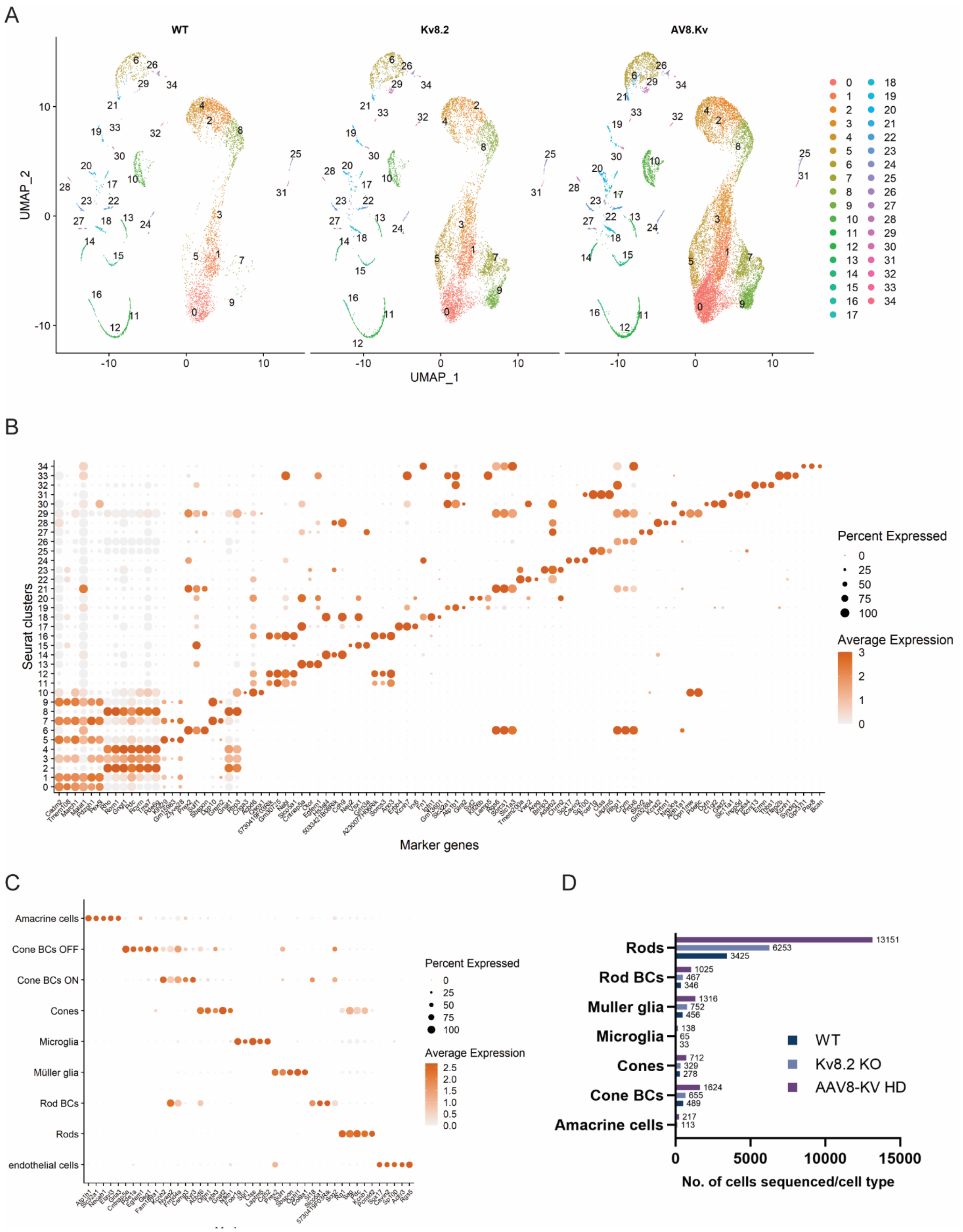


**Supplementary Figure 2. Additional single-cell RNA sequencing data analysis of retinal cell populations. (A)** UMAP visualization of retinal cells from wild-type (WT), Kv8.2 knockout (KO), and AAV8-Kv8.2 treated groups, showing distinct clustering of major cell types. **(B)** Heatmap of marker gene expression across identified clusters, highlighting cell-type-specific markers such as *Cadm2*, *Tmem108*, *Malat1*, and *Gad2*. **(C)** Bar plot summarizing the number of cells sequenced per cell type, including rods, cones, Müller glia, rod bipolar cells (BCs), cone BCs (ON/OFF), amacrine cells, and microglia. **(D)** Dot plot of average expression and percent expression for selected marker genes across major retinal cell types.

***Supplementary Table 1. Number of cells analysed for each group and cell type***

| Cell type | Wild type | Kv8.2 KO*[1] | AAV8.KCNV2*[2] | hKNCV2*[3] | Relative % |
|---|---|---|---|---|---|
| **Amacrine cells** | 51 | 113 | 215 | 2 | 0.93 |
| **Cone BCs** | 489 | 655 | 1613 | 11 | 0.07 |
| **Cones** | 278 | 329 | 535 | 180 | 33.6 |
| **Microglia** | 33 | 65 | 134 | 4 | 2.98 |
| **Muller glia** | 456 | 752 | 1250 | 66 | 5.28 |
| **Rod BCs** | 346 | 467 | 1019 | 6 | 0.58 |
| **Rods** | 3425 | 6253 | 9929 | 3222 | 32.4 |

*BCs, Bipolar cells.*

*[1] Number of cells sequenced within each cell group from Kv8.2 KO untreated retinas*

*[2] Number of cells sequenced within each cell group from Kv8.2 KO treated retinas*

*[3] Number of cells within each cell group from Kv8.2 KO treated retinas expressing the human KCNV2 gene after treatment*
